# Horizontal and vertical transmission of transgenerational memories via the *Cer1* transposon

**DOI:** 10.1101/2020.12.28.424563

**Authors:** Rebecca S. Moore, Rachel Kaletsky, Chen Lesnik, Vanessa Cota, Edith Blackman, Lance R. Parsons, Zemer Gitai, Coleen T. Murphy

**Author notes:** Equal contribution (Rebecca S. Moore, Rachel Kaletsky).

## Abstract

Animals face both external and internal dangers: pathogens threaten from the environment, and unstable genomic elements threaten from within. Previously, we discovered that *C. elegans* protects itself from pathogens by “reading” bacterial small RNAs and using this information to both induce avoidance and transmit memories for several generations. Here we found that these memories can be transferred to naïve animals via *Cer1* retrotransposon-encoded capsids. *Cer1* functions at the step of transmission of information from the germline to neurons, and is required for *C. elegans’* learned avoidance ability and for mothers to pass this information on to progeny. The presence of the *Cer1* retrotransposon in wild *C. elegans* strains correlates with the ability to learn and inherit small RNA-induced pathogen avoidance. Together, these results suggest that *C. elegans* has co-opted a potentially dangerous retrotransposon to instead protect itself and its progeny from a common pathogen through its inter-tissue signaling ability, hijacking this genomic element for its own adaptive immunity benefit.

## Introduction

The transmission of information across generations through non-genetic means, or transgenerational epigenetic inheritance (TEI), was long thought to be impossible due to the Weismann barrier between the germline and somatic cells, which preserves immortal germ cells in their pristine state. However, recent data from worms (Houri-Zeevi et al., 2020; Rechavi et al., 2014; Webster et al., 2018), flies (Bozler et al., 2019), and mice (Dias and Ressler, 2014) suggest that inheritance of stress responses may help animals survive in harsh environments. We recently discovered that *C. elegans* passes small RNA-mediated learned *Pseudomonas aeruginosa* avoidance behavior on to several generations of progeny through a molecular mechanism that requires an intact germline and neuronal signaling (Kaletsky et al., 2020a). This process requires uptake of a *P. aeruginosa* small RNA called P11, processing through the RNA interference pathway, piRNA regulation and P granule function in the germline, downregulation of a neuronal gene with complementarity to a specific bacterial small RNA, and gene expression changes in the ASI sensory neuron (Kaletsky et al., 2020a). This small RNA-mediated process enables mothers and four generations of her progeny to avoid pathogenic *Pseudomonas aeruginosa*.

The question of whether animals can transmit memories to one another has a storied and controversial history (McConnell et al., 1959; Shomrat and Levin, 2013), but recent work in *Aplysia* suggests that RNA from the CNS of trained animals can induce a form of non-associative long-term memory when injected into naïve animals (Bédécarrats et al., 2018). Whether these horizontally transferred memories could be transmitted transgenerationally, thereby breaking the Weismann barrier, or in a natural context, has not yet been addressed.

Here, we find that whole-worm lysates from the grandprogeny of trained *C. elegans* can transmit memory of learned avoidance and transgenerational inheritance of that avoidance behavior to naïve animals and their four generations of progeny, through virus-like particles encoded by the *Cer1* retrotransposon. In addition to its role in horizontal memory transfer, *Cer1* is required within individuals for small-RNA mediated learned pathogen avoidance and transgenerational epigenetic inheritance through its ability to convey information from the germline to neurons. *Cer1’s* presence in the genomes of wild strains correlates with their ability to carry out these behaviors, a beneficial role for *Cer1* in contrast to its previously reported deleterious effects (Dennis et al., 2012). Thus, *Cer1* function may provide *C. elegans* long-lasting protection from pathogens in their natural environments.

## Results

### Transgenerational memories are horizontally transferred to naïve worms

*C. elegans* is initially attracted to *Pseudomonas aeruginosa*, but learns to avoid this pathogen after exposure (Zhang et al., 2005) (Figure 1A-B). Worms learn to avoid *P. aeruginosa* (PA14) through several independent mechanisms involving bacterial small RNAs, metabolites, and additional pathogenesis factors (Kaletsky et al., 2020a; Meisel et al., 2014; Singh and Aballay, 2019; Troemel et al., 2006) (Figure S1). However, small RNA-mediated learned avoidance is the only pathway that leads to transgenerational memory inheritance (Kaletsky et al., 2020a) (Figure S1). To test whether transgenerational learned avoidance can be horizontally transferred to naïve worms, we used mechanical homogenization to prepare crude lysates from wild-type grand-progeny (F2) of P11- or control-trained grandmothers (Figure 1C-D, Figure S2A-B). Naïve animals were exposed to the lysate on *E. coli* plates for 24h, then tested for *P. aeruginosa* avoidance learning. We found that lysate from F2s of P11-trained, but not control-trained grandmothers, was sufficient to induce naïve worms to avoid *P. aeruginosa* (Figure 1D-E), indicating horizontal transmission of memory.

**Figure 1.**
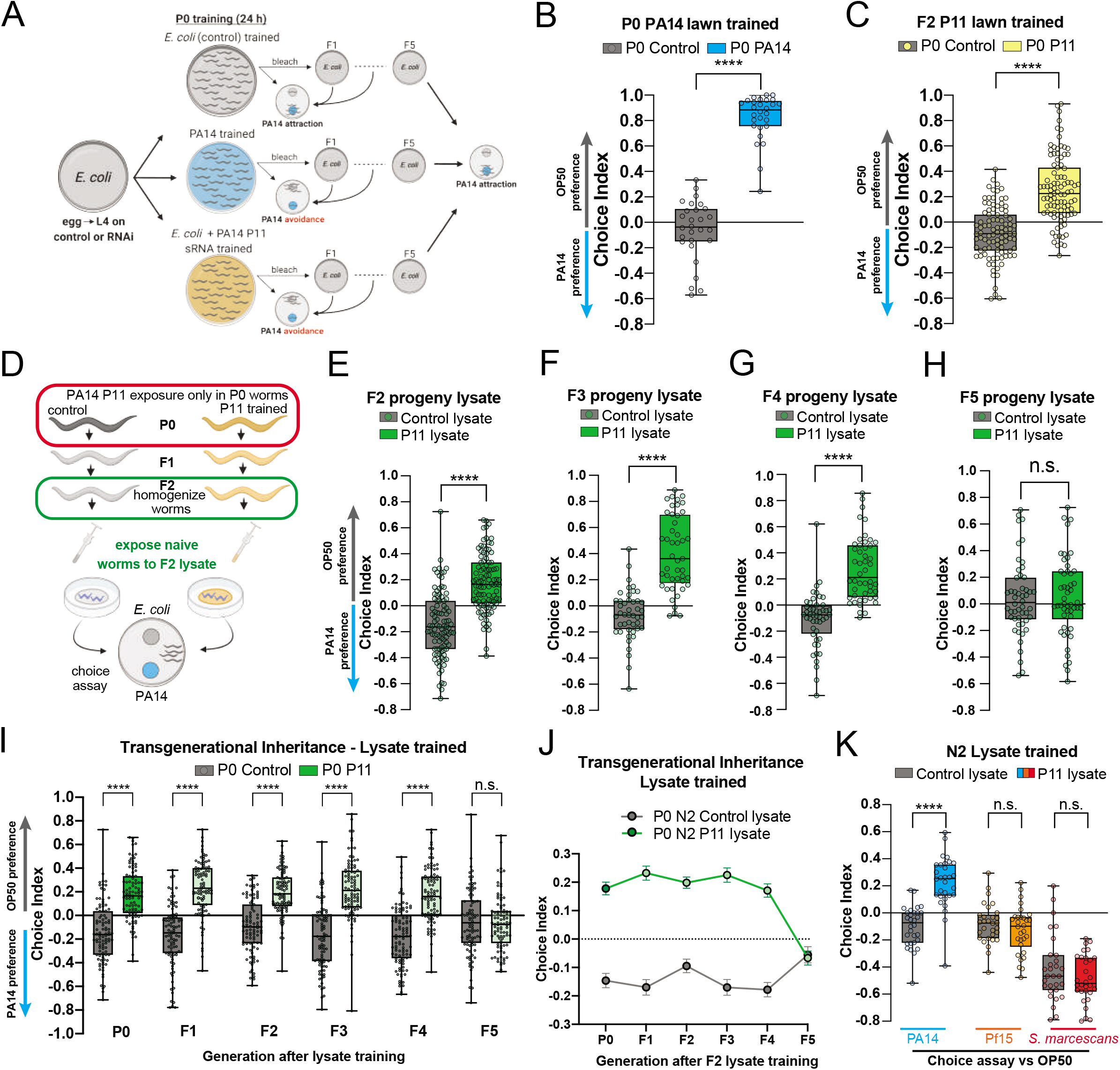
Horizontal transmission of transgenerational PA14 avoidance learning. A, Worms were trained on non-pathogenic OP50, PA14 lawns, or *E. coli* expressing the PA14 P11 small RNA or a control. Choice assays to OP50 versus PA14 bacteria were then performed. Trained animals were bleached to maintain subsequent generations without additional PA14 or P11 exposure. Choice index - (number of worms on OP50 - number of worms on PA14)/(total number of worms). B, Worms exposed to a PA14 lawn (for 24 h) learn to avoid PA14. C, F2 progeny of P11-trained grandmothers inherit PA14 avoidance behavior compared to controls. D, Schematic of protocol for horizontal memory transfer. F2 progeny from control or P11-trained grandmothers are homogenized, and naïve worms are exposed to the F2 lysate for 24 h before testing avoidance behavior. E-H, PA14 avoidance behavior in naïve animals trained with worm lysate from F2s (E), F3s (F), F4s (G), or F5 animals (H) derived from P0-control or P11 trained mothers. F2 thorough F4 worm lysates confer PA14 avoidance (E-G), while F5 worm lysate does not (H). I-J, The avoidance behavior acquired by naïve worms trained with lysate from F2 progeny from P11-trained grandmothers (compared to controls) is inherited in progeny through the F4 generation. K, Naïve worms were trained with lysate from F2s grand-progeny of control or P11 −trained grandmothers, as in D. After lysate exposure, worms were split into groups and tested in 3 different choice assays: OP50 v. PA14 (left), OP50 vs Pf15 (middle), or OP50 vs *S. marcescens* (right). Each dot represents an individual choice assay plate (average of 115 worms per plate) from all replicates. At least 3 biological replicates for all experiments. Box plots: center line, median; box range, 25–75th percentiles; whiskers denote minimum-maximum values. Unpaired, two-tailed student’s t-test (B-H), One-way analysis of variance (ANOVA) (I), Tukey’s multiple comparison test, \*\*\*\**P* < 0.0001, NS, not significant. Supplementary Table 1 for exact sample sizes (*n*) and *P* values.

We previously observed that training of mothers with either *P. aeruginosa* or P11 small RNA induces a memory of learned avoidance that lasts through the F4 generation (Kaletsky et al., 2020a; Moore et al., 2019). While the lysate from F2-F4 progeny can induce learning in naïve animals, lysate from the F5 generation - which does not show inheritance of learned behavior from either *P. aeruginosa* or P11 training - is not able to transfer learned avoidance (Figure 1E-H, Figure S2C-E). Furthermore, progeny of lysate-trained P0 animals inherited this learned avoidance behavior, lasting through the F4 generation after training (Figure 1I-J), indicating that transgenerational inheritance can occur after horizontal transfer of memory.

We also previously established that training animals on *P. aeruginosa* or P11 small RNA induces avoidance specifically against *P. aeruginosa*, rather than to other bacteria (Kaletsky et al., 2020b; Moore et al., 2019). To test whether the horizontally-acquired memory is specific to *P. aeruginosa*, lysate-trained animals were tested for changes in preference to *Pseudomonas fluorescens* (Pf15) or *Serratia marcescens*. While worms exposed to lysate from grandprogeny of P11-trained grandmothers learned to avoid *P. aeruginosa* compared to controls, lysate training did not alter the worms’ attraction to either *P. fluorescens* (Pf15) or *S. marcescens* (Figure 1K). These results indicate that the horizontally-transferred memories are specifically encoded for *P. aeruginosa* avoidance and are likely not caused by a non-specific response that induces broad neuronal changes in preference.

### Virus-like particles in lysate transmit transgenerational memories to naïve worms

RNA has been implicated in the transfer of memory from the CNS of trained *Aplysia* to naïve animals (Bédécarrats et al., 2018), and we previously showed that the mechanism by which *C. elegans* learns the identity of pathogenic *Pseudomonas* requires bacterial small RNAs (Kaletsky et al., 2020a). Therefore, we tested 1) whether free, total RNA isolated from F2s of trained animals could transfer memory, and 2) whether the trained F2 lysate would still transfer memory if treated with RNase before worm training. However, total RNA from trained F2s was not able to induce avoidance learning (Figure 2A), and RNase treatment of the trained F2 lysate did not abolish memory transfer (Figure 2B).

**Figure 2.**
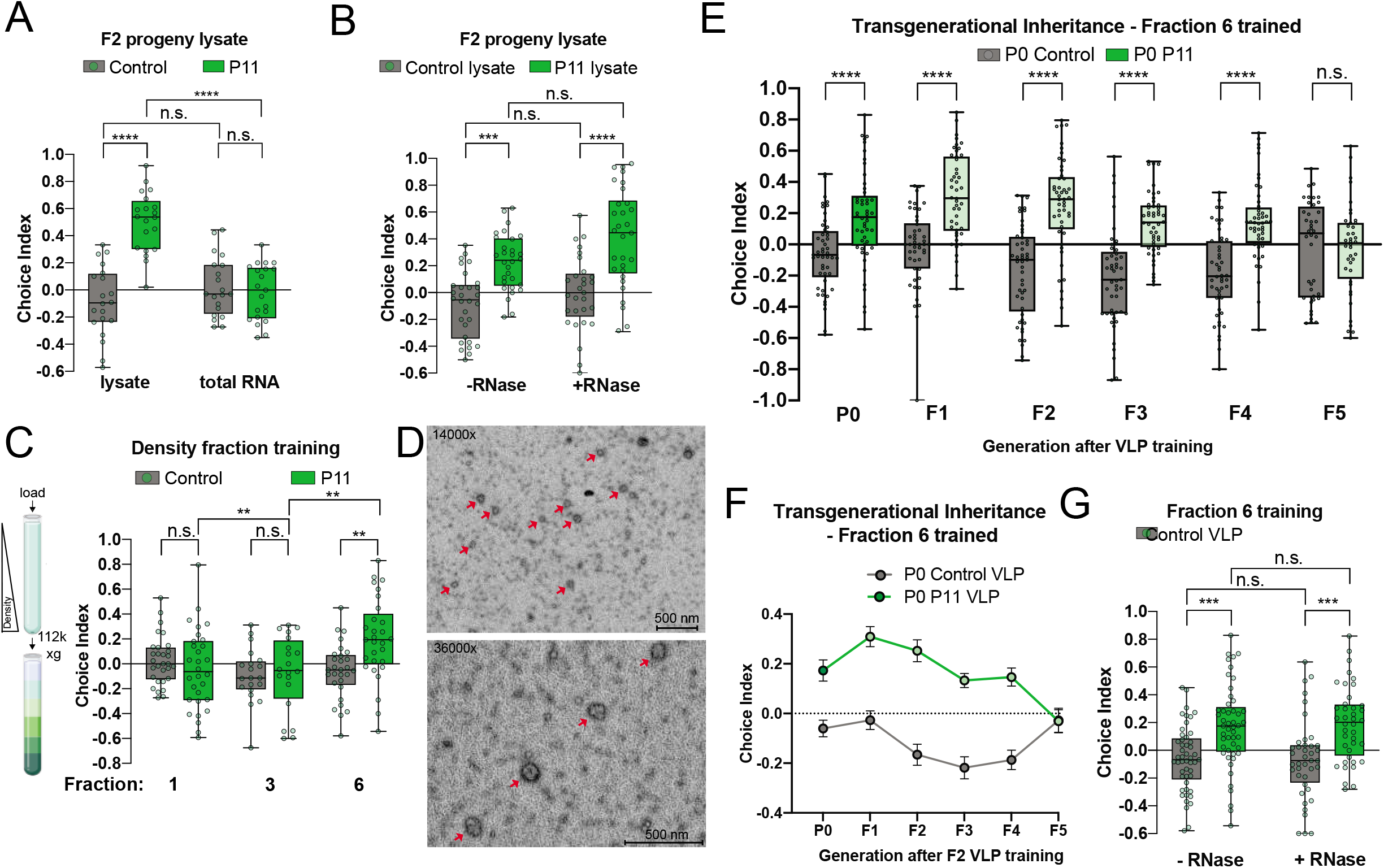
Transgenerational memories are transferred to naïve worms via virus-like particles. A, Training with total RNA isolated from F2 progeny descended from control or P11-trained grandmothers does not confer PA14 avoidance learning compared to F2 lysate training. B, Memory conferred by F2 worm lysate training is resistant to RNAse treatement. Lysate was exposed to RNAse prior to naïve worm exposure. C, F2 worm lysates were fractionated using density-based centrifugation. Fractions 1, 3, and 6 from the gradient were used to train naïve worms, followed by PA14 choice assays. D, Negative-stain electron microscopy was performed on fraction 6. E-F, Transgenerational inheritance of PA14 avoidance in progeny of worms exposed to fraction 6 (derived from F2s from control P11-trained grandmothers). G, PA14 avoidance behavior induced by fraction 6 is resistant to RNAse treatment. Fraction 6 was exposed to RNAse prior to naïve worm exposure. Each dot represents an individual choice assay plate (average of 115 worms per plate) from all replicates. At least 3 biological replicates for all experiments. Box plots: center line, median; box range, 25-75th percentiles; whiskers denote minimum-maximum values. One-way (E) or two-way (A-C, G) analysis of variance (ANOVA), Tukey’s multiple comparison test. *\*\*P* ≤ 0.01, ***P ≤ 0.001, **”*P* < 0.0001, NS, not significant. Supplementary Table 1 for exact sample sizes (n) and *P* values.

While these results would suggest RNA is not involved, another possibility is that the information could be protected; for example, a similar RNase treatment of Arc virus-like particles (VLPs) still allows transfer of Arc mRNA between neurons (Pastuzyn et al., 2018). To determine if purified capsids might carry the memory of P11 training, we tested density-fractionated lysates from F2s of P11-trained grandmothers for their ability to induce avoidance. Only the densest fraction (#6), which should contain VLPs, induced *P. aeruginosa* avoidance behavior in naïve animals (Figure 2C).

To determine whether capsids or virus-like particles were present in the fraction that induced learning behavior, we performed electron microscopy on the densest fraction (Figure 2D) and identified ~90 nm virus-like particles (VLPs). This fraction was also able to induce behavior not only in the trained generation, but also through the F4 generation (Figure 2E-F). Although there was too little material to successfully build a library for RNA-seq, the Bioanalyzer trace of the RNase-treated VLPs shows that there is RNA inside the capsids (Figure S3A), and RNase treatment of the VLP fraction did not prevent the induction of avoidance learning (Figure 2G), supporting the model that capsids protect cargo RNA.

### *Cer1* is required for small RNA-induced pathogen avoidance learning and transgenerational memory

The capsids we observed by EM were similar in size to VLPs made by the *Cer1* retrotransposon (Dennis et al., 2012). *Cer1* has homology to the Ty3/Gypsy retrotransposon (Figure 3A), and forms VLPs that are detectable by EM and present in the germline of N2 animals at 20°C (Figure 3B). Therefore, we investigated whether *Cer1* might be involved in learned pathogen avoidance and its inheritance. The *Cer1* GAG protein was detected in the densest fraction (#6), which induced learned avoidance (Figure 2C) in wild-type worms (Figure 3C-D). A point mutation (G6369A) in *Cer1* abolishes its detection by immunofluorescence (Figure 3C) or by Western blot (Figure 3D), suggesting that this mutation prevents expression of *Cer1* gene products. *Cer1#* mutant mothers were still able to learn on a *P. aeruginosa* lawn (Figure 3E), consistent with intact routes of lawn learning, such as innate immunity and metabolites; however, loss of *Cer1* abolishes the F1 inheritance of *P. aeruginosa* avoidance behavior (Figure 3F), which functions through the separate small RNA-mediated pathway (Kaletsky et al., 2020a). Reduction of *Cer1* via RNAi also abrogated *P. aeruginosa-mediated* pathogen avoidance inheritance (Figure 3G-H). Loss of *Cer1* by mutation or RNAi also completely abrogated the ability of mothers trained on *E. coli*+P11 to learn *P. aeruginosa* avoidance (Figure 3I-J). Unlike *Cer1*, loss of a different Ty3/Gypsy retrotransposon, *Cer4*, had no effect on learning or transgenerational memory induced by PA14 lawn or *E. coli*+P11 training of N2 mothers (Figure S4).

**Figure 3.**
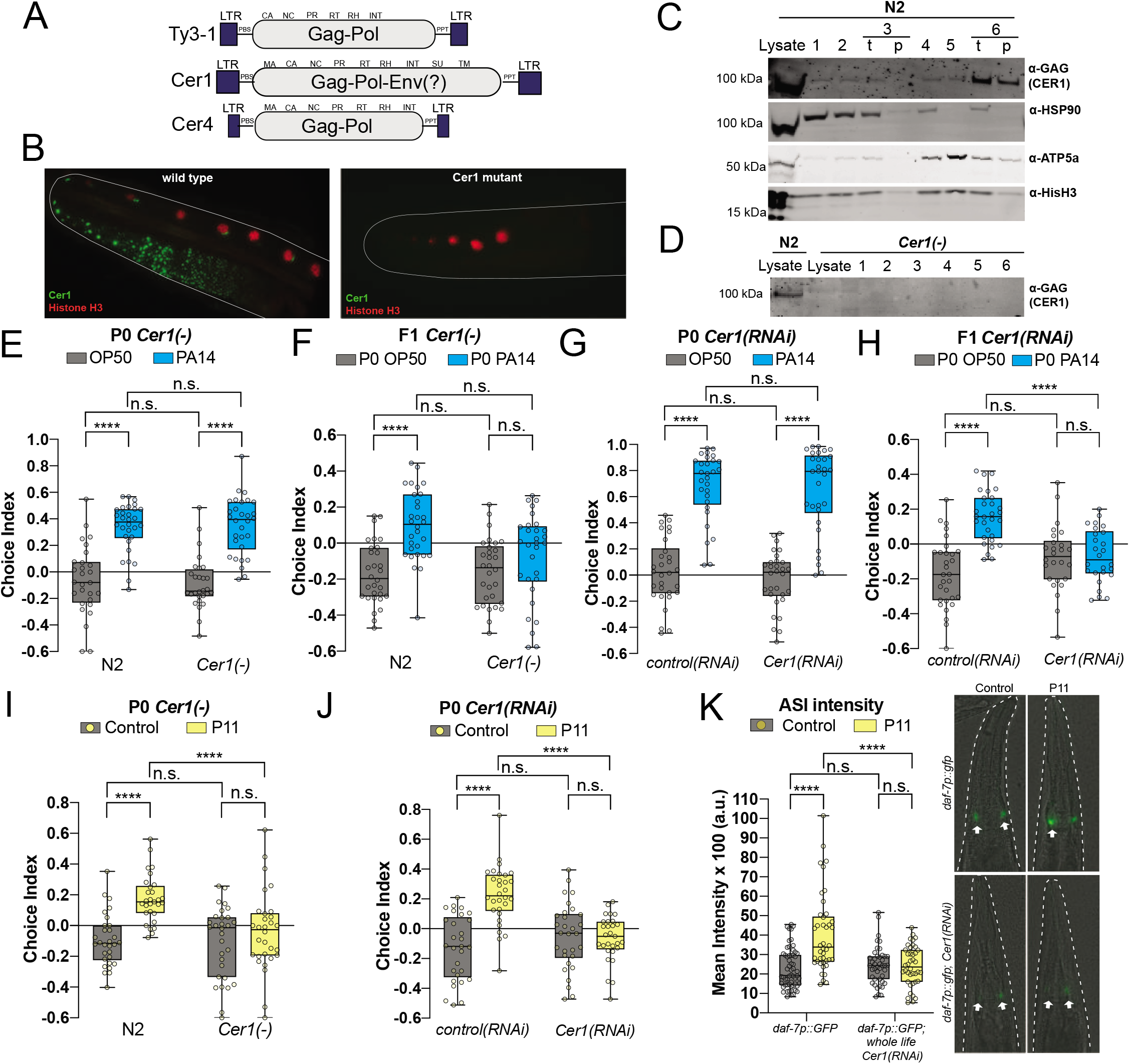
*Cer1* is required for P11-acquired learning and vertically inherited memory. A, Schematic of *C. elegans Cer1, Cer4*, and *S. cerevisiae* Ty3-1. (LTR = Long Terminal Repeat, PBS = Primer Binding Site, MA = Matrix, CA = Capsid, NC = Nucleocapsid, PR = Protease, RT = Reverse Transcriptase, RH = RNaseH, INT = Integrase, SU = Surface, TM = Transmembrane, PPT = PolyPurine Tract. B, Immunofluorescence of wild type or *Cer1* mutant worm germlines stained for Histone H3 (control) or *Cer1* GAG. C, *Cer1* GAG is detected prominently in fraction 6 by western blot, while other cellular markers are enriched in lighter fractions. D, Western blot of *Cer1* GAG in wild type and *Cer1* mutant animals. GAG is absent from all fractions in the *Cer1*(*gk870313*) mutants (G6369A substitution). E-F, *Cer1*(*gk870313*) mutants have a normal naive preference for PA14, and can learn to avoid PA14 bacteria lawns (E), but are defective for transgenerational inheritance of avoidance behavior (F). G-H, Similar to *Cer1* mutants, worms treated with whole-life *Cer1* RNAi can learn to avoid PA14 bacterial lawns (G), but are defective for transgenerational inheritance of avoidance behavior (H). I, *Cer1* mutant worms cannot learn to avoid PA14 through P11 small RNA exposure. J, P11-induced learning is defective in worms treated with whole-life *Cer1* RNAi. K, *daf-7p::gfp* expression in ASI neurons (white arrows) increases upon P11 small RNA exposure compared to controls, *daf-7p::gfp* expression does not increase upon P11 exposure in *Cer1* RNAi-treated worms. Scale bar, 25 μm. Each dot represents an individual choice assay plate (average of 115 worms per plate) from all replicates. At least 3 biological replicates for all experiments. Box plots: center line, median; box range, 25-75th percentiles; whiskers denote minimum-maximum values. Two-way (E-K) analysis of variance (ANOVA), Tukey’s multiple comparison test, \*\*\*\**P* < 0.0001, NS, not significant. Supplementary Table 1 for exact sample sizes (*n*) and *P* values.

Upon training with *P. aeruginosa* or P11 small RNA, *daf-7p::gfp* expression increases in the ASI sensory neuron (Kaletsky et al., 2020a; Meisel et al., 2014; Moore et al., 2019). Loss of *Cer1* prevents this increase in expression, suggesting that *Cer1* acts upstream of the regulation of *daf-7* expression in the ASI neuron (Figure 3K). Together, these results suggest that *Cer1* is required for small RNA-mediated pathogen avoidance in mothers and their progeny, is present in the VLP fraction that induces learning, and acts upstream of neurons in the small RNA-mediated learning pathway. To determine whether Cer1 is required for not only vertical memory transmission to progeny, but also for horizonal memory transfer, we prepared worm lysates and VLP-containing fractions from wild type and *Cer1* mutant F2s from control or P11-trained grandmothers. Consistent with the requirement for *Cer1* in horizontal memory acquisition, neither the lysate nor the analogous density-purified fraction (fraction #6) isolated from *Cer1* mutants derived from P11-trained grandmothers were able to induce avoidance of *P. aeruginosa* (Figure 4A-B). These results suggest that *Cer1* capsids are required for the horizontal transfer of transgenerational epigenetic memories to naïve worms.

**Figure 4.**
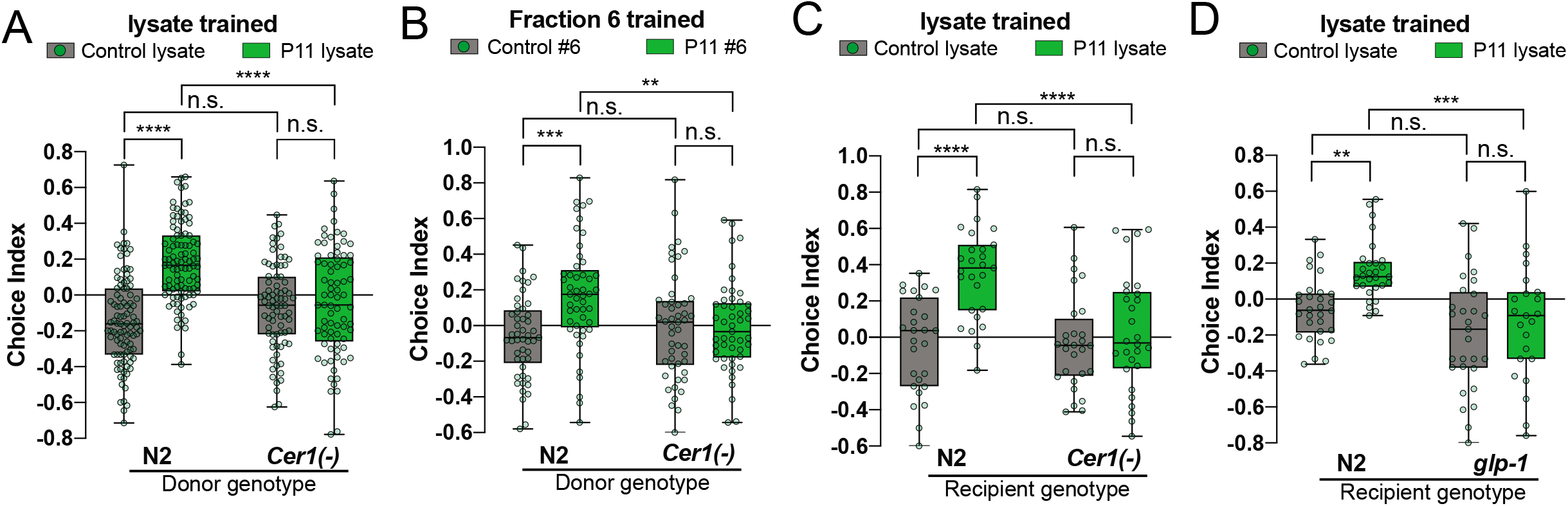
Cer1 is required for horizontal transmission of learned avoidance. A-B, F2 lysate (A) or virus-like particles (B) from *Cer1* mutant worms does not induce horizontal memory transfer compared to wild type F2 lysate. Each F2 worm lysate (wild type or *Cer1* mutant) were the grand-progeny from control or P11 −trained grandmothers. Lysate from wild-type or *Cer1* mutant F2 was used to train naïve wild-type animals. C-D, Wild-type F2 worm lysate was obtained from the grand-progeny of control or P11-trained grandmothers and used to train naïve recipient *Cer1* mutants (C) or germline-less *glp-1* worms (D) compared to wild-type recipient controls. Each dot represents an individual choice assay plate (average of 115 worms per plate) from all replicates. At least 3 biological replicates for all experiments. Box plots: center line, median; box range, 25–75th percentiles; whiskers denote minimum-maximum values. Two-way (E-M) analysis of variance (ANOVA), Tukey’s multiple comparison test. \*\**P* ≤0.01, \*\*\**P* ≤ 0.001, \*\*\*\**P* < 0.0001, NS, not significant. Supplementary Table 1 for exact sample sizes (*n*) and *P* values.

Since *Cer1* is required for both vertical and horizontal transfer of pathogen avoidance learning, we next asked whether *Cer1* and/or a germline is required in recipient worms, or if treating with *Cer1-* containing lysate bypasses the requirement for *Cer1* in recipient animals (for example, by direct uptake by neurons). *Cer1* mutants trained with wild-type F2 lysates were unable to learn *P. aeruginosa* avoidance (Figure 4C). Germline-less *glp-1*(*e2141*) also failed to learn *P. aeruginosa* avoidance upon F2 lysate training (Figure 4D). These results show that both *Cer1* and a functional germline are required in recipient animals for horizontal memory transfer through Cer1 capsid.

### *Cer1* is required for transmission of germline state of avoidance learning

Our results show that *Cer1* is required in mothers for small RNA-mediated learned avoidance and in their progeny for the inheritance of this behavior. Previously, we found that the process of inducing transgenerational inheritance of pathogen avoidance requires uptake of small, non-coding RNAs from *Pseudomonas*, processing of this small RNA in the intestine and germline, and transmission of an unknown signal that is conveyed to the ASI neurons to influence avoidance behavior (Kaletsky et al., 2020a).

To determine the mechanism of *Cer1*’s function in learned pathogen avoidance and its inheritance, we wanted to determine the step at which it is required - the initiation of the transgenerational signal, maintenance of this signal in the germline from generation to generation, or a subsequent, post-germline step that results in execution of avoidance behavior (transmission of the signal from germline to neurons or neuronal function) (Figure 5A). The step at which *Cer1* acts in the pathway was not clear from our experiments, because a mutant or *Cer1* RNAi for several generations would not distinguish a lasting and permanent effect of *Cer1* activation from a transient effect that only affects one step of the transgenerational learned pathogen avoidance process. However, these steps can be distinguished through a simple experiment: knockdown of the gene of interest in the F1 generation after P0 training, followed by control RNAi in generations F2-F5. Knockdown of a gene involved in initiation (P0) would have no effect if reduced only in the F1 generation (Figure 5A, blue line, “initiation”); F1 knockdown of a gene involved in germline maintenance or propagation would permanently eliminate learned behavior (orange line, “maintenance/ propagation”); and F1 knockdown of a gene that only functions in transmission of the signal or functions in neurons would eliminate the behavior for a generation or two, but should return once the RNAi knockdown is ended (green line, “behavior”).

**Figure 5.**
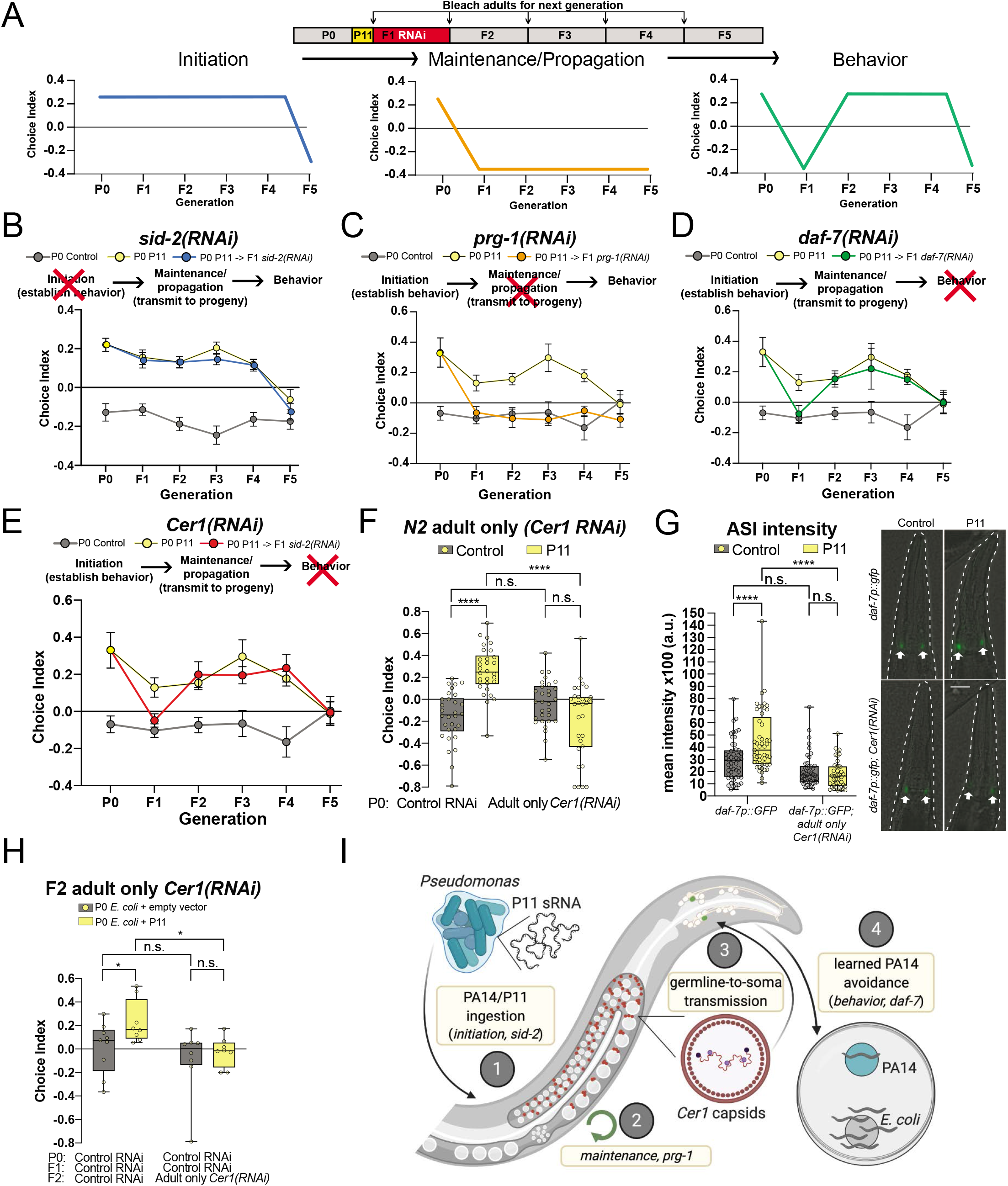
*Cer1* is required for execution of PA14 small RNA-mediated transgenerational inheritance of avoidance behavior. A, Schematic of F1 RNAi treatment following control or P11 exposure in PO mothers. Reducing F1 expression of a gene required for initiation of transgenerational inheritance should have no effect on behavior (blue), while reduced F1 expression of a TEI maintenance gene should eliminate memory in the F1 and subsequent generation (orange). F1 knockdown of a gene required for the execution of behavior should affect F1 behavior, but not that of subsequent generations (green). B-E, Wild-type mothers were trained with control or P11-expressing *E. coli*. F1 progeny were then treated with either *sid-2* (B), *prg-1* (C), *daf-7* (D), or *Cer1* (E) RNAi. Subsequent generations of progeny were maintained on normal food and examined for PA14 avoidance behavior. F, Worms were treated with *Cer1* RNAi only during adulthood (from L4 to Day 1) before training on control or P11-expressing *E. coli. Cer1* is required specifically during adulthood for learned avoidance behavior in mothers. G, *daf-7p::gfp* expression does not increase upon P11 exposure in *Cer1* adult-only RNAi-treated worms. Scale bar, 25 μm. H, F2 progeny of control or P11 grandmothers were treated with *Cer1* adult-only RNAi. *Cer1* is required in adults in F2 worms that have inherited PA14 avoidance behavior. I, Model of germline-to-soma communication of PA14 avoidance through *Cer1*. Each dot represents an individual choice assay plate (average of 115 worms per plate) from all replicates. At least 3 biological replicates for all experiments. Box plots: center line, median; box range, 25-75th percentiles; whiskers denote minimum-maximum values. Two-way (F-G) analysis of variance (ANOVA), Tukey’s multiple comparison test. * *P* < 0.05, \*\*\*\**P* < 0.0001, NS, not significant. Supplementary Table 1 for exact sample sizes (*n*) and *P* values.

Knockdown of *sid-2*, the “NA transporter that is expressed in the intestine (McEwan et al., 2012), only in F1 does not affect behavior in any generation, likely because its role is to facilitate uptake of bacterial small RNAs from the gut, which is critical in initiation (P0) but is not needed in later generations (Figure 5B, Figure S5A). By contrast, knockdown of the piRNA Piwi/Argonaute PRG-1 in the F1 generation eliminates behavior not only in F1, but also causes a permanent loss of avoidance behavior (Figure 5C, Figure S5B). These results are consistent with previous data suggesting that *prg-1* is required for maintenance or propagation of avoidance behavior, and that loss of *prg-1* erases transgenerational memory (Ashe et al., 2012). The TGF-beta ligand DAF-7 is expressed in the ASI neuron, and is required to execute the avoidance behavior (Kaletsky et al., 2020a; Moore et al., 2019). Reduction of *daf-7* by RNAi in the F1 generation following maternal *P. aeruginosa* (Figure S5C) or *E. coli*+P11 (Figure 5D) training abrogated avoidance behavior in the same generation (F1). However, progeny raised on control RNAi recovered their avoidance behavior in the F2-F4 generations (Figure 5D, Figure S5C), demonstrating that the encoded memory was retained even when *daf-7* expression was reduced, and that avoidance behavior could return. This shows that *daf-7* is not required for germline maintenance of transgenerational memory, but is instead involved in the execution of avoidance behavior. These *sid-2, prg-1*, and *daf-7* RNAi initiation vs maintenance vs execution behavior results, respectively, agree with their previously-determined roles in intestine (McEwan et al., 2012), germline (Batista et al., 2008), and neurons (Kaletsky et al., 2020a; Moore et al., 2019; Ren et al., 1996).

*Cer1* capsids are present in the germline, and their presence depends on *prg-1* and P granules in worms (Dennis et al., 2012); in yeast, Ty3 VLP formation is similarly dependent on P-bodies (Beliakova-Bethell, 2006). Therefore, we first hypothesized that *Cer1* might function at the step of maintaining the transgenerational signal in the germline, similar to *prg-1* (Figure 5C). However, while *Cer1*(*RNAi*) treatment in the F1 progeny of wild-type mothers trained with *E. coli* expressing P11 (Figure 5E, S5E, right) or with *P. aeruginosa* (Figure S5D, left) caused loss of avoidance behavior, the avoidance memory recovered in subsequent generations maintained on control RNAi allowing *Cer1* re-expression (F2-F4; Figure 5E). These results resembled *daf-7* knockdown and recovery, rather than the permanent loss of learned avoidance that *prg-1* knockdown causes, suggesting that *Cer1* acts in the execution of avoidance behavior rather than at the step of maintenance of the transgenerational signal. This further suggested that *Cer1*’s role in learned pathogen avoidance might not be restricted to germline function, despite the fact that it is primarily expressed in the germline (Dennis et al., 2012), but rather may act at a step between germline and neuron function.

To test the notion that *Cer1* might act in a post-germline, dynamic, transient step, we carried out RNAi starting in adulthood. First, knockdown of *Cer1* in trained P0 adults (Figure 5F) blocked avoidance learning as well as whole-life RNAi treatment did (Figure 3J), showing that *Cer1* can be knocked down effectively in adults. Similarly, loss of Cer1 only in adults prevents the induction of *daf-7p::gfp* expression in the ASI (Figure 5G). Knockdown of *Cer1* in trained P0 adults followed by treatment on control RNAi in F1 allowed the re-emergence of avoidance behavior (Figure S6A-D), further establishing that *Cer1* is not involved in establishment of the transgenerational signal. Knockdown of *Cer1* only in adults of the F2 generation abrogated behavior (Figure 5H, Figure S6E), despite the F1 animals having demonstrated inheritance of avoidance (Figure S6C-D). Together, these results suggest that the process is dynamic: if the transgenerational inheritance of avoidance had been set by regulation of neuronal gene expression levels in the embryonic state, then knockdown of *Cer1* should not have affected behavior. Instead, we see that *Cer1*, which acts upstream of *daf-7* in the ASI, dynamically regulates behavior in adult animals.

Together, these results show that loss of *Cer1* does not erase transgenerational memory, but rather is required downstream of the memory maintenance machinery in order to execute avoidance behavior. Thus, its role is unlikely to be solely in the germline, but more likely in the communication of the status of avoidance state information from the germline to the neurons in every generation. This germline-to-soma signaling (Figure 5I) ultimately affects neuronal activity and behavior to avoid a common pathogen, and also improves their survival on that pathogen (Moore et al., 2019). Together, these functions might provide an evolutionary benefit from the insertion and activity of a retrotransposon that was previously thought to be solely deleterious.

### The ability of wild strains of *C. elegans* to carry out small RNA-induced pathogen avoidance learning and transgenerational memory correlates with *Cer1* expression

Roughly 15% of the *C. elegans* genome consists of transposon genetic material (Laricchia et al., 2017). The Ty3/Gypsy family retrotransposon *Cer1* is one of these elements, and is inserted into the genomes of roughly 70% of wild *C. elegans* strains (Palopoli et al., 2008), although the sites of these insertions differ - some are present in the *plg-1* locus, which regulates “plugging” upon mating, while others are present elsewhere (Laricchia et al., 2017) (Figure 6A). Similarly, some *Cer1* insertions are only remnants of the active transposon, with only LTRs (long terminal repeats) detectable (Figure 6A). Therefore, we wondered whether the presence of full-length *Cer1* in the genomes of strains isolated from the wild is necessary or sufficient to confer the ability to learn and remember pathogen avoidance.

**Figure 6.**
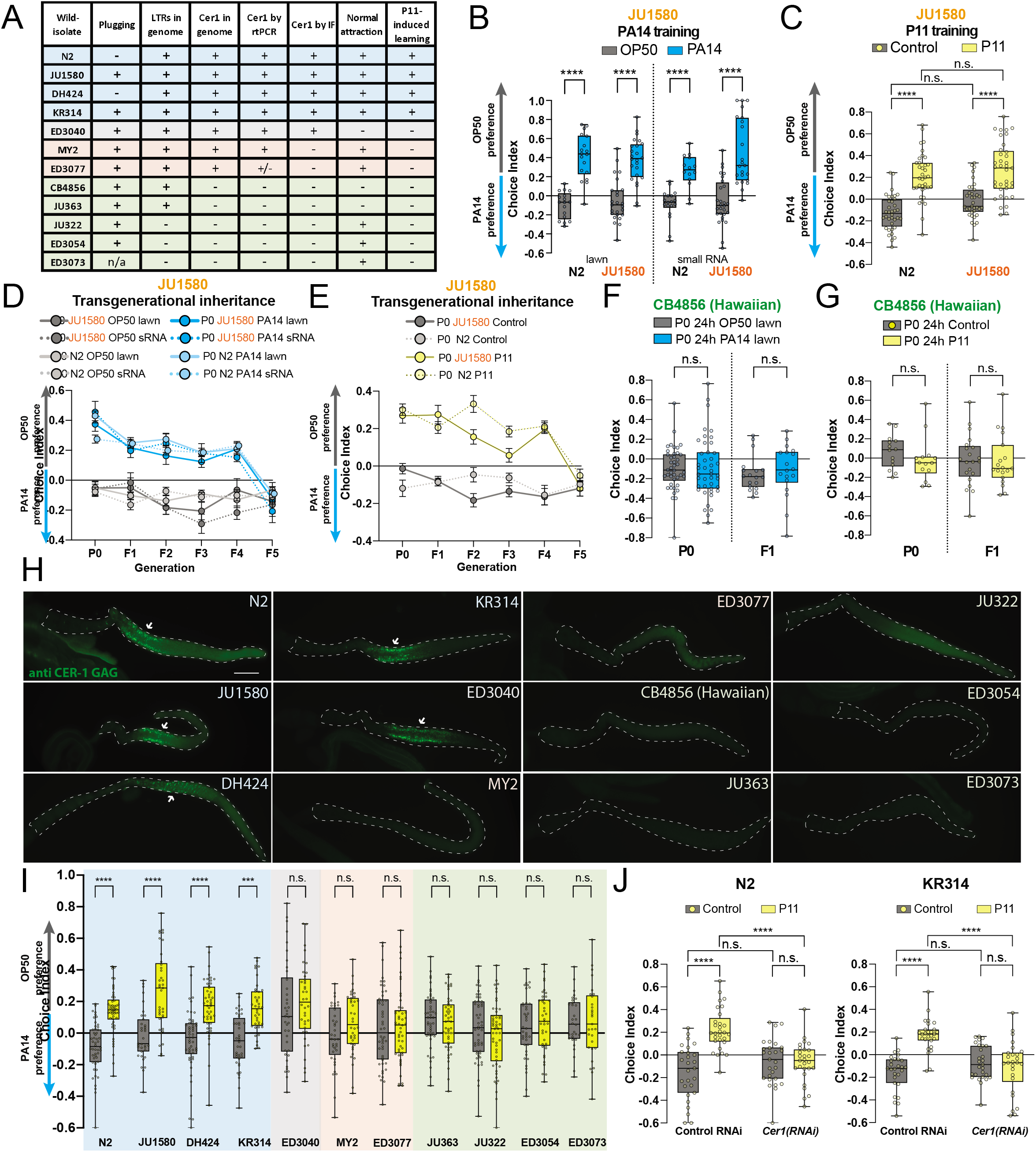
*Cer1* expression correlates with PA14 avoidance learning in *C. elegans* wild isolates. A, *C. elegans* wild isolates were characterized for plugging(Palopoli et al., 2008), presence and expression of Cer1, naïve PA14 attraction, and P11 small RNA-induced learning. B, *C. elegans* wild-isolate JU1580 mothers exposed to PA14 lawns (left) or small RNAs (right) learn to avoid PA14 in a choice assay. C. JU1580 mothers exposed to *E. coli* expressing P11 learn to avoid PA14 after training compared to controls. D, Like wild-type N2 *C. elegans*, progeny of JU1580 PA14 lawn- or small RNA-trained mothers continue to avoid PA14 through the F4 generation. 5^th^ generation progeny return to naïve preference. E, Progeny of JU1580 *E. coli*+P11-trained mothers continue to avoid PA14 for four generations (F1-F4) before attraction to PA14 resumes in the 5^th^ generation. F-G, *C. elegans* Hawaiian mothers exposed to PA14 bacteria lawns (F) or *E.* coli+*P11* (G) do not learn to avoid PA14, and progeny of trained mothers do not inherit avoidance behavior. H, Immunofluorescence of *Cer1* GAG was visualized in *C. elegans* wild isolates. I, PA14 avoidance behavior in wild isolate mothers trained on control bacteria or P11-expressing *E. coli*. J, Whole-life RNAi knockdown of *Cer1* in N2 (left) and KR314 (right) eliminates P11-induced PA14 learned avoidance. Each dot represents an individual choice assay plate (average of 115 worms per plate) from all replicates. At least 3 biological replicates for all experiments. Box plots: center line, median; box range, 25–75th percentiles; whiskers denote minimum–maximum values. One-way (B, F, G) or two-way (C, J) analysis of variance (ANOVA), Tukey’s multiple comparison test, \*\*\**P* ≤ 0.001, \*\*\*\**P* < 0.0001, NS, not significant. Supplementary Table 1 for exact sample sizes (*n*) and *P* values.

An intact copy of *Cer1* is present in the wild strain JU1580, as shown by the complete coverage of the coding sequences and LTRs by de novo assembly (Cook et al., 2017) (Figure S7A). We found that like N2, JU1580 animals learn to avoid *P. aeruginosa* both through exposure to the pathogen (Figure 6B, left) and small RNAs (Figure 6B, right), as well as by exposure to *E. coli*+P11 (P11 training) (Figure 6C). Furthermore, trained JU1580 mothers can pass this information on to their progeny for four generations, just as N2 does (Figure 6D-E). These results suggest that the mechanisms underlying transgenerational inheritance of learned pathogen avoidance via small RNAs are conserved.

In contrast to our findings with JU1580, another *C. elegans* strain, CB4856 (“Hawaiian”), is unable to learn to avoid *P. aeruginosa* after lawn (Figure 6F) or *E. coli*+P11 training (Figure 6G), or to pass this information on to its progeny (F1). It was previously shown that Hawaiian does not have *Cer1* inserted into its genome (Palopoli et al., 2008) (Figure S7C), but this is not the only difference between N2 and Hawaiian. CB4856 and N2 differentially survive on *P. aeruginosa*, and this difference is mediated by the *npr-1* gene, which regulates leaving behavior in response to oxygen levels. However, the genomic region of *npr-1* in JU1580 has the “wild” SNP of *npr-1*, as Hawaiian does (Cook et al., 2017), ruling out *npr-1* as the source of the difference in pathogenic learning ability. Similarly, the *maco-1* gene, which is downregulated upon exposure to *P. aeruginosa* and is required for learned *P. aeruginosa* avoidance behavior (Kaletsky et al., 2020a), is identical between N2 and Hawaiian in the 17 nucleotides of homology to P11 (Figure S7B), suggesting that Hawaiian’s inability to learn and pass on learned avoidance is not due to a lack of sequence matching between P11 small RNA and its *maco-1* target.

To determine whether the presence of *Cer1* correlates with the ability to learn pathogen avoidance more widely in nature, we examined the expression of *Cer1* RNA via RT-PCR (Figure 6A) and the presence of *Cer1* GAG protein via immunofluorescence (Figure 6H) in N2, JU1580, Hawaiian, and an additional nine wild strains of *C. elegans*, and we tested the ability of these wild strains to carry out P11-mediated learned avoidance of *P. aeruginosa*. Like N2 and JU1580, the wild strains DH424 and K4314 expressed *Cer1* RNA and Cer1 GAG protein, and were able to learn *P. aeruginosa* avoidance after P11 training (Figure 6A, H-I). Other strains behaved like Hawaiian, as they were unable to learn P11-induced avoidance and were defective for attraction to *P. aeruginosa* (Figures S6); none of these strains had *Cer1* inserted into the genome or expressed *Cer1* at appreciable levels (Figure 6A, H). (Although the twelfth strain, ED3040, has *Cer1* inserted into its genome and expresses *Cer1*, it is defective for normal attraction to *P. aeruginosa* and does not exhibit increased avoidance upon training.) Finally, treatment of the *Cer1*-expressing wild strain KR314 with *Cer1* RNAi abolished its P11-mediated learning (Figure 6J). Thus, the presence and expression of *Cer1* in wild strains of *C. elegans* largely correlates with ability to learn to avoid *P. aeruginosa* after small RNA-mediated training.

## Discussion

Here we have shown that information conveying pathogenic exposure status can be transferred from trained to naïve *C. elegans*, via capsids of Ty3/Gypsy *Cer1* retrotransposon. Additionally, the transfer of this information induces memory that lasts for four additional generations, similar to training on *Pseudomonas aeruginosa* or its small RNA, P11. Our results provide a molecular mechanism by which memory transmission might occur: the *Cer1* retrotransposon expresses virus-like particles that can confer memory of learned pathogen avoidance to other individuals, and within an individual, from germline to neurons. Thus, memories of learned avoidance of pathogens can be transferred between individuals, and can induce transgenerational inheritance of the learned information.

The idea that memory can be transferred between individuals is old but controversial. Reports of horizontally-transferred memory in planarians (McConnell et al., 1959) seemed to contradict both the concept of memory storage occurring only at synapses and the strict protection of the germline from somatic changes proposed by Weismann in the late 1800s. These findings were more recently supported by an independent study in Planaria that used an automated system to reduce bias (Shomrat and Levin, 2013). However, planaria divide asexually, and thus the concept of a Weismann barrier might be less strict. Furthermore, no molecular mechanism for this type of memory transfer has been determined. Another example of memory transfer between individuals is from recent work in *Aplysia*, in which the RNA extracted from the CNS of trained animals injected into naïve animals was able to increase sensitization in a DNA methylation-dependent manner (Bédécarrats et al., 2018), an example of an epigenetic mechanism of memory storage, but whether this could happen in the wild or influence the behavior of progeny is unknown. Our results in *C. elegans* suggest that the *Cer1* retrotransposon enables the transfer of a memory of a pathogen from germline to nervous system, between generations, and from animal to animal.

The fact that *Cer1*’s presence in wild strains of *C. elegans* correlates with the ability to learn and transgenerationally inherit pathogen avoidance suggests that *Cer1* itself may have enabled the acquisition of this behavior. *C. elegans* dies within 23 days in the presence of *Pseudomonas aeruginosa*, killing mothers before they have finished reproducing, which would deleteriously affect their fitness. *Cer1* was previously noted to reduce fecundity in non-pathogenic conditions (Dennis et al., 2012), but here we found that the presence of *Cer1* enables the worms to learn to avoid *Pseudomonas*. If naïve animals are able to take up *Cer1* capsids from animals who have died and lysed, it would allow them to acquire learned avoidance without experiencing illness themselves (Figure 7), effectively vaccinating them against future *P. aeruginosa* exposure by inducing avoidance behavior. Furthermore, as infected mothers often “bag” (die of matricide), the ability of other worms to take up *Cer1* VLPs might provide them with the ability to avoid the pathogen - perhaps the first utilization of memory transfer. The ability to avoid pathogens for multiple generations could provide *C. elegans* that have acquired *Cer1* an advantage in environments rife with pathogens.

**Figure 7.**
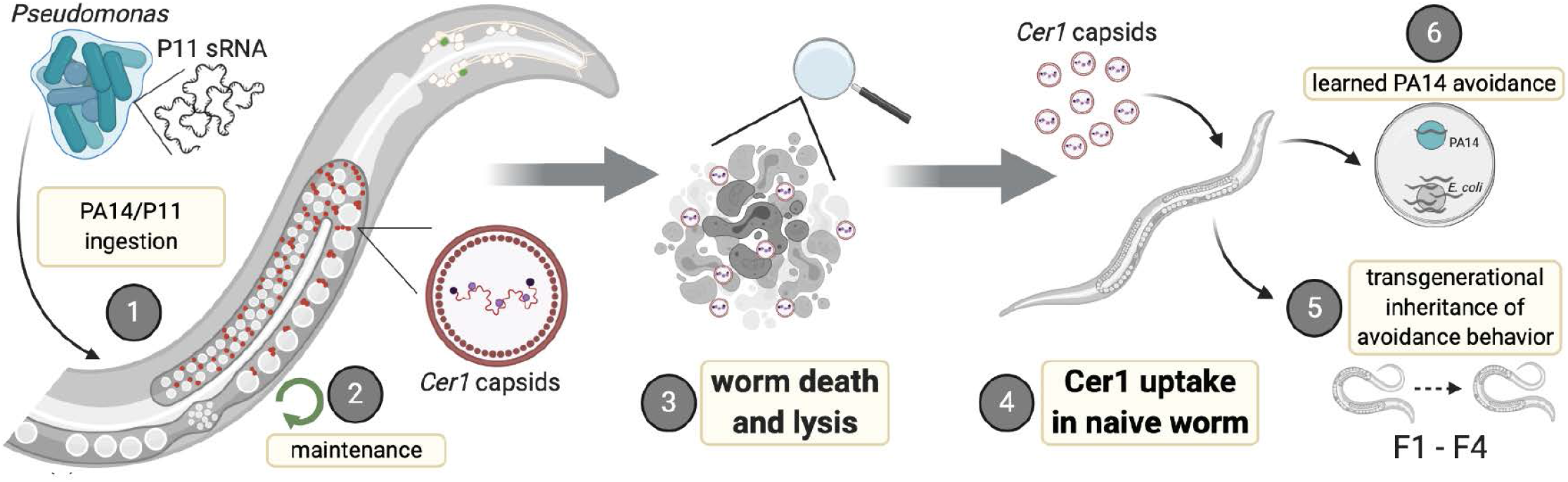
Model of horizonal memory transfer vai *Cer1*. Horizontal transfer of PA14 avoidance memory occurs when naïve worms are exposed to *Cer1*’s virus-like particles from an animal that has already inherited the memory. Uptake of *Cer1* induces memory directly in that animals and in four generations of its progeny.

Here we have shown that rather than being solely deleterious (Dennis et al., 2012), the presence of the *Cer1* retrotransposon in fact may have been co-opted by *C. elegans* to help it survive in an environment that requires frequent encounters with pathogens. The ability of the *Cer1* retrotransposon to confer a benefit to the host is surprising considering the classical nature of transposons in genomes. Transposons are highly abundant in animal genomes, and generally regarded as pernicious, mutagenic genetic elements whose mobility can lead to disease and the erosion of host fitness. Transposons incur damage to hosts on several fronts: through misregulation of host processes, such as interfering with host transcription, processing of mRNAs, and chromatin structure (Elbarbary et al., 2016), or through disruption of the host genome through transposition. Consistent with other transposons, the presence of *Cer1* was previously only noted to be deleterious, as its expression decreases fecundity and lifespan in non-pathogenic conditions (Dennis et al., 2012). The finding that *Cer1* is required for learned and transgenerationally-inherited PA14 avoidance behavior shows that ancient retrotransposons can be co-opted and repurposed to benefit the worm, an example of transposon-host mutualism (Feschotte and Gilbert, 2012). Since retrotransposition in *C. elegans* has never observed under laboratory conditions (Bessereau, 2006; Laricchia et al., 2017), it is likely that *Cer1* mediates this acquired worm behavior independent of its potential for novel genome insertion as a retrotransposon.

While the domestication of transposons underlies some of the most critical transitions in animal evolution (Agrawal et al., 1998; Dupressoir et al., 2012; Hiom et al., 1998; Sheen and Levis, 1994; Smit and Riggs, 1996; Tudor et al., 1992) the requirement for *Cer1* in transgenerational learned behavior is unique in that *Cer1* is an active transposon, and that *Cer1* confers a behavioral ability, avoidance, on the animals. An interesting parallel arises with comparison to recent studies of Arc (of Ty3/Gypsy family origin), which showed that Arc VLPs can transmit cellular genetic material across neurons in a process that underlies synaptic plasticity in fly and mammalian brains (Ashley et al., 2018; Lyford et al., 1995; Pastuzyn et al., 2018). While *C. elegans* lacks a direct *Arc* ortholog, *Cer1* is also a member of the Ty3/Gypsy family and similarly forms capsids (Dennis et al., 2012). *Cer1*’s role in pathogen avoidance, and specifically in the avoidance behavior step - rather than in generation or maintenance of the transgenerational memory - was surprising, given the fact that *Cer1* produces VLPs in the germline; however, VLPs are also present in non-germline cells at lower abundance, perhaps suggesting at least a transient presence outside of the germline (Dennis et al., 2012). Although it is possible that *Cer1* acts like Arc, transmitting information between neurons, a more parsimonious explanation, given the abundance of *Cer1* VLPs in the germline and our genetic evidence placing it upstream of *daf-7* regulation in the ASI neuron, is that germline *Cer1* VLPs carry host cargo to neurons, where subsequent changes in expression and activity modulate behavior (Fig 5C).

Our data suggest that *Cer1* functions in a novel, dynamic germline-to-neuron signaling mechanism that may represent the co-option of retrotransposon function to improve *C. elegans’* survival, and its progeny’s survival, in pathogenic environments. *Cer1* appears to provide *C. elegans* immediate protection from abundant pathogenic *Pseudomonas* species in its environment, but also confers lasting generational benefits by communicating an adaptive immune signal of learned avoidance to its descendants. Moreover, the ability to provide memories of pathogen avoidance to neighboring worms might allow greater survival of its kin.

## Supporting information

Statistics Reporting

## Acknowledgements

We thank the *C. elegans* Genetics Center for strains; the EM facility at the Princeton Imaging and Analysis Center; J. Priess for providing anti-*Cer1* GAG antibody hybridoma; S. Petry for equipment; BioRender for model figure design software; J. Shepherd and the laboratory of C.T.M. for discussion; and R. Clausen for assistance. C.T.M. is the Director of the Glenn Center for Aging Research at Princeton and an HHMI-Simons Faculty Scholar. This work was supported by the Glenn Foundation for Medical Research (GMFR CNV1001899), the HHMI-Simons Faculty Scholar Program (AWD1005048), and a Pioneer Award to C.T.M. (NIGMS DP1GM119167), T32GM007388 (NIGMS) support of R.S.M., and a Pioneer Award to Z.G. (DP1A1124669).

---

**Supplemental Figure 1.**
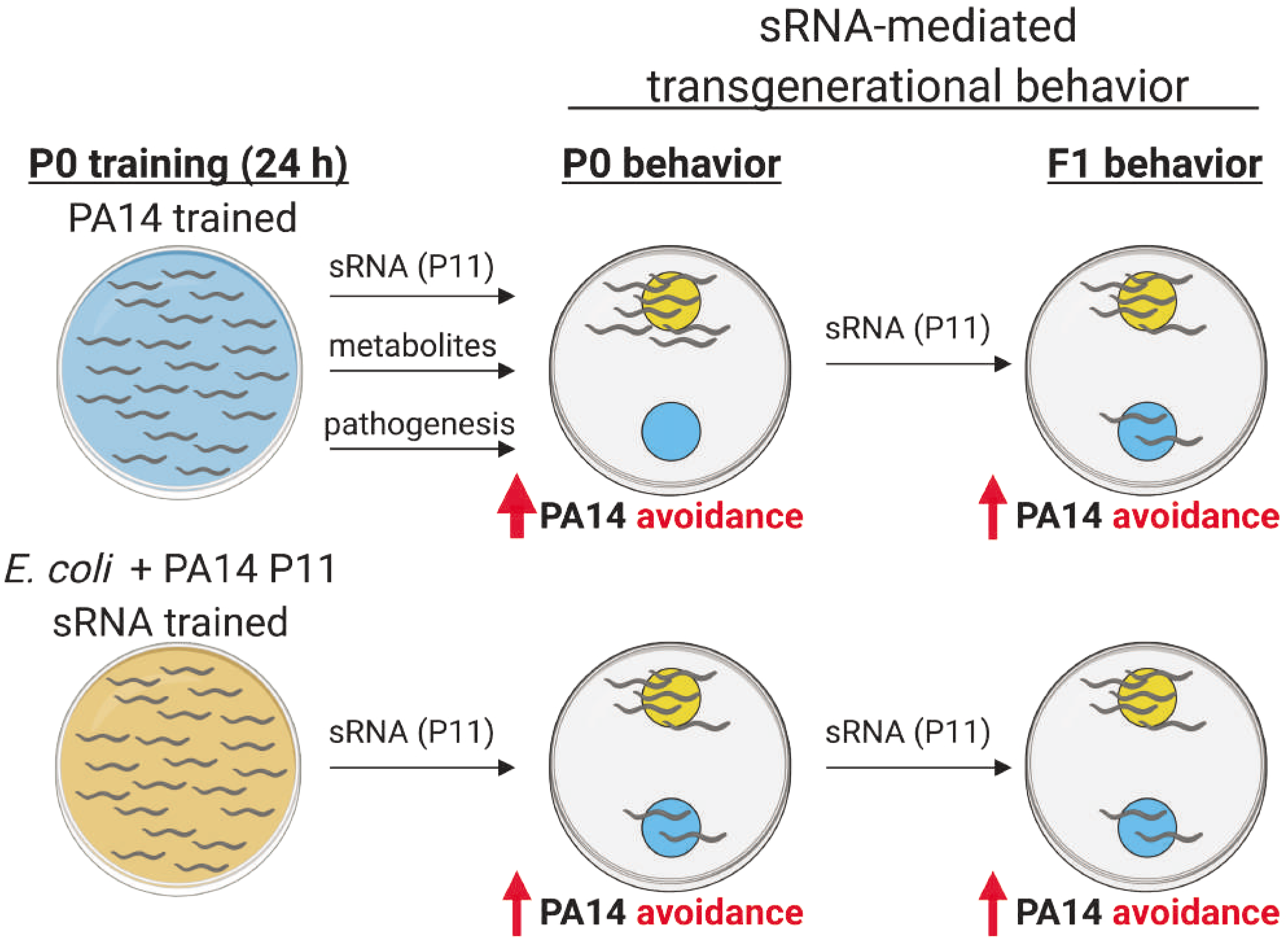
Modes of PA14 and P11-induced learning and inheritance. (A) Naïve *C. elegans* prefer PA14 if given a choice between OP50 (*E. coli*) and PA14. After exposure to PA14 for 24h, worms learn to avoid PA14 via three cues: (1) small RNAs (specifically P11), (2) metabolites and, (3) innate immune pathways. This avoidance behavior can be transgenerationally inherited in naive progeny for four generations, before resetting in the 5th. Only small RNAs are required for transgenerational inheritance of pathogen avoidance.

**Supplemental Figure 2.**
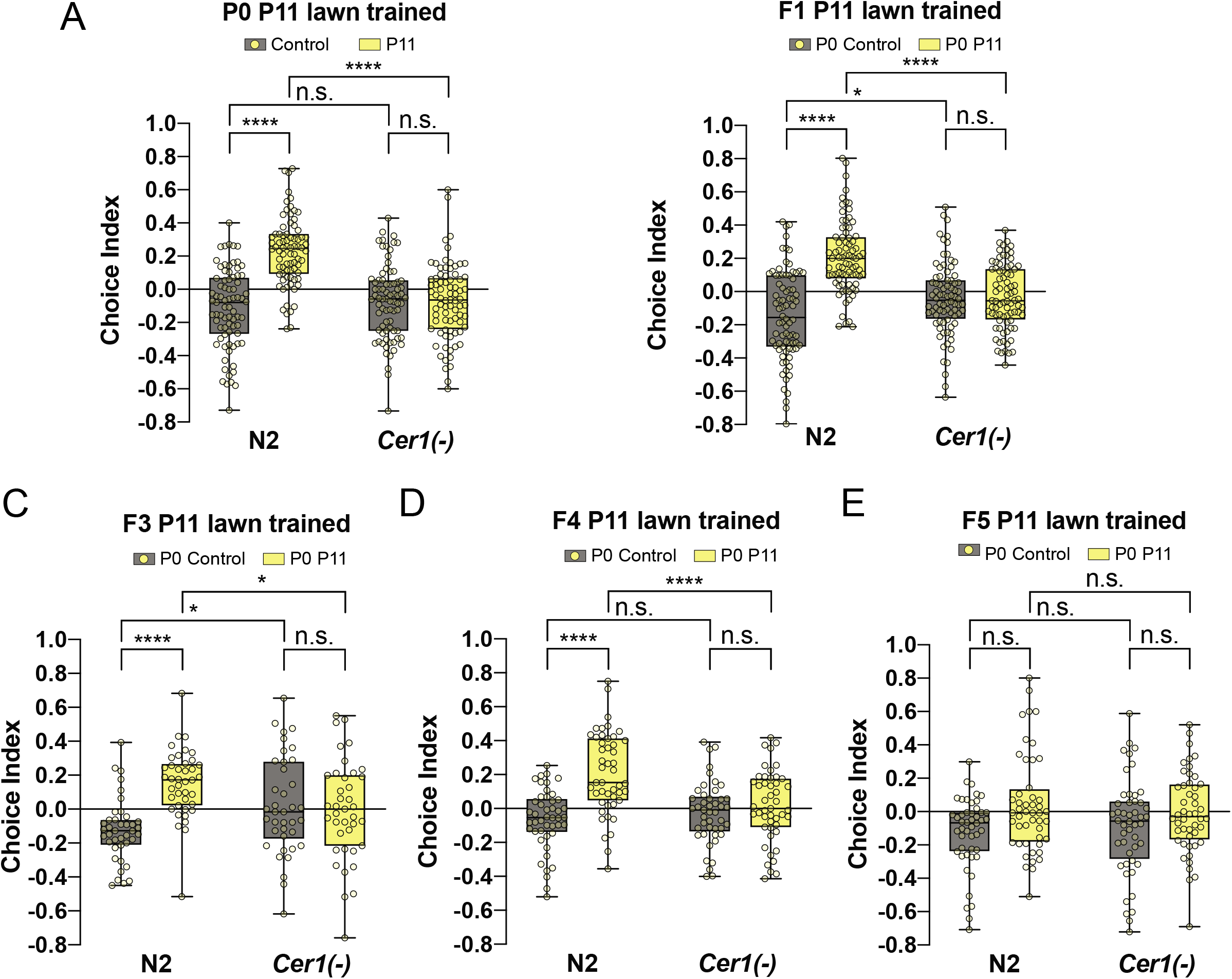
Learned and inherited P11-induced PA14 avoidance behavior in donor worms used for lysate training of naïve animals. A, Mothers trained on P11-expressing *E. coli* learn to avoid PA14 compared to controls. F1 (B), F2 (Figure 1C), F3 (C), and F4 (D) progeny inherit PA14 avoidance memory from their ancestors. E, PA14 avoidance memory is reset in the F5 generation. A-D, *Cer1* mutant mothers cannot learn to avoid PA14 upon P11 exposure, and the F1-F4 progeny (B-D) also do not exhibit transgenerational memory. Each dot represents an individual choice assay plate (average of 115 worms per plate) from all replicates. At least 3 biological replicates for all experiments. Box plots: center line, median; box range, 25-75th percentiles; whiskers denote minimum-maximum values. Two-way (A-E) analysis of variance (ANOVA), Tukey’s multiple comparison test. * *P* < 0.05, \*\*\*\**P* < 0.0001, NS, not significant. Supplementary Table 1 for exact sample sizes (*n*) and *P* values.

**Supplemental Figure 3.**
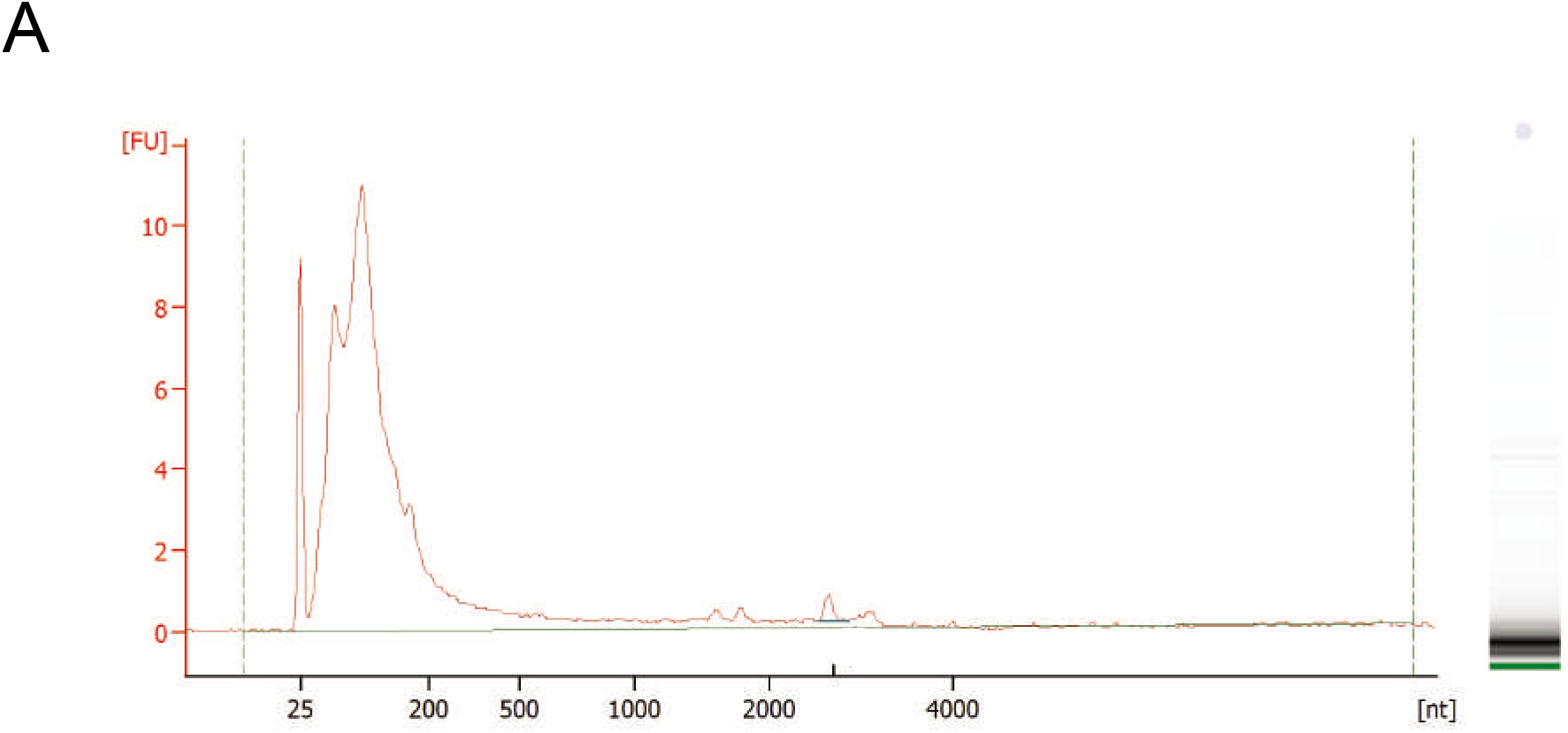
RNA profile from isolates VLP fraction. Total RNA was extracted from VLPs (fraction 6) following RNAse-treatment and subsequent RNase inactivation (Figure 2G). RNA was analyzed using a Bioanalyzer RNA 600 Pico kit.

**Supplemental Figure 4.**
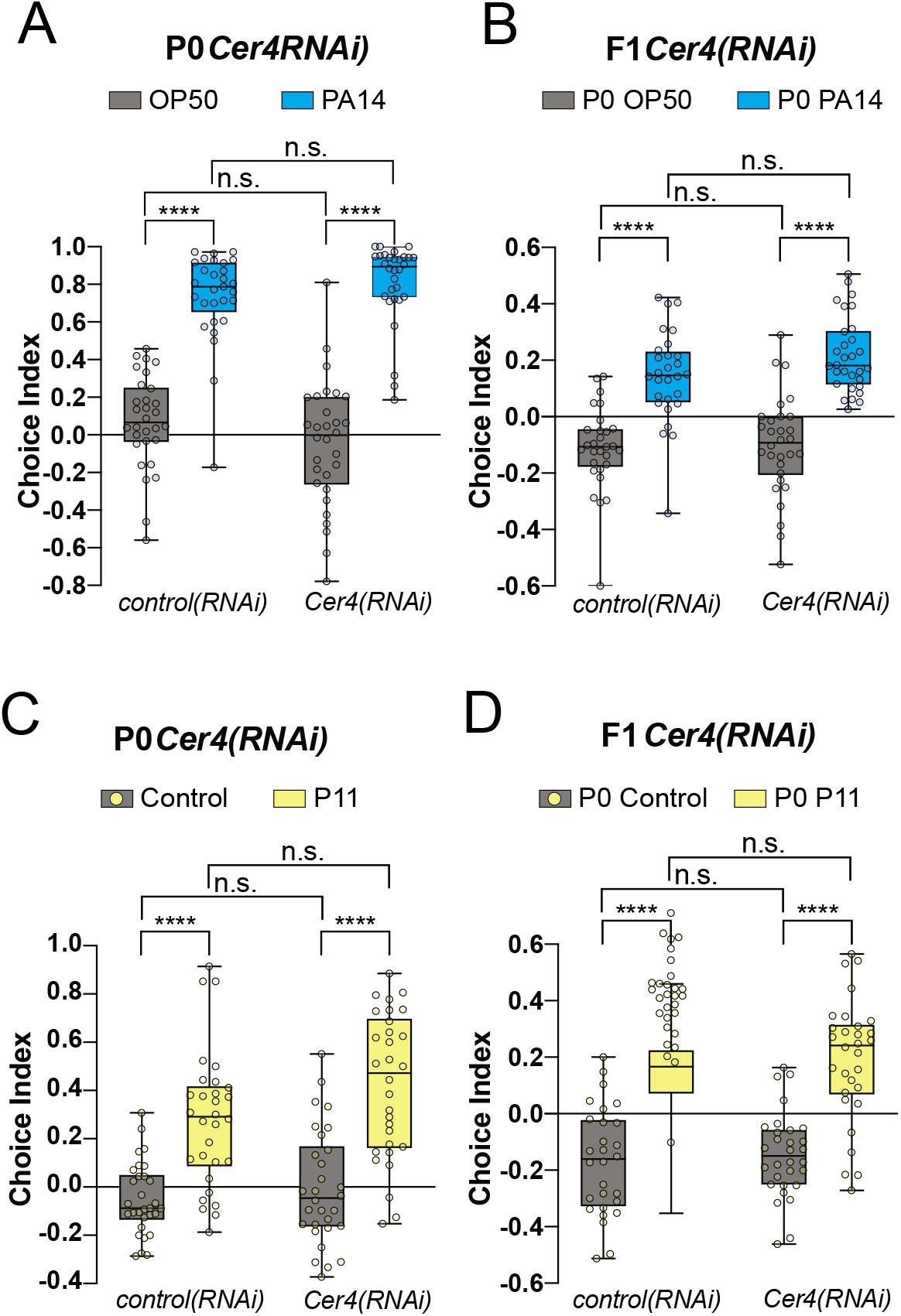
PA14 avoidance behavior in worms with Cer4-RNAi knockdown. A-B, *Cer4* is not required for PA14 lawn-induced learning (A) or transgenerational memory inheritance (B). C-D, P11 small RNA-induced learning (C) and transgenerational memory (D) is intact in worms treated with *Cer4* RNAi. Each dot represents an individual choice assay plate (average of 115 worms per plate) from all replicates. At least 3 biological replicates for all experiments. Box plots: center line, median; box range, 25-75th percentiles; whiskers denote minimum-maximum values. Two-way (A-D) analysis of variance (ANOVA), Tukey’s multiple ∞mparison test, \*\*\*\**P* < 0.0001, NS, not significant. Supplementary Table 1 for exact sample sizes (*n*) and *P* values.

**Supplemental Figure 5.**
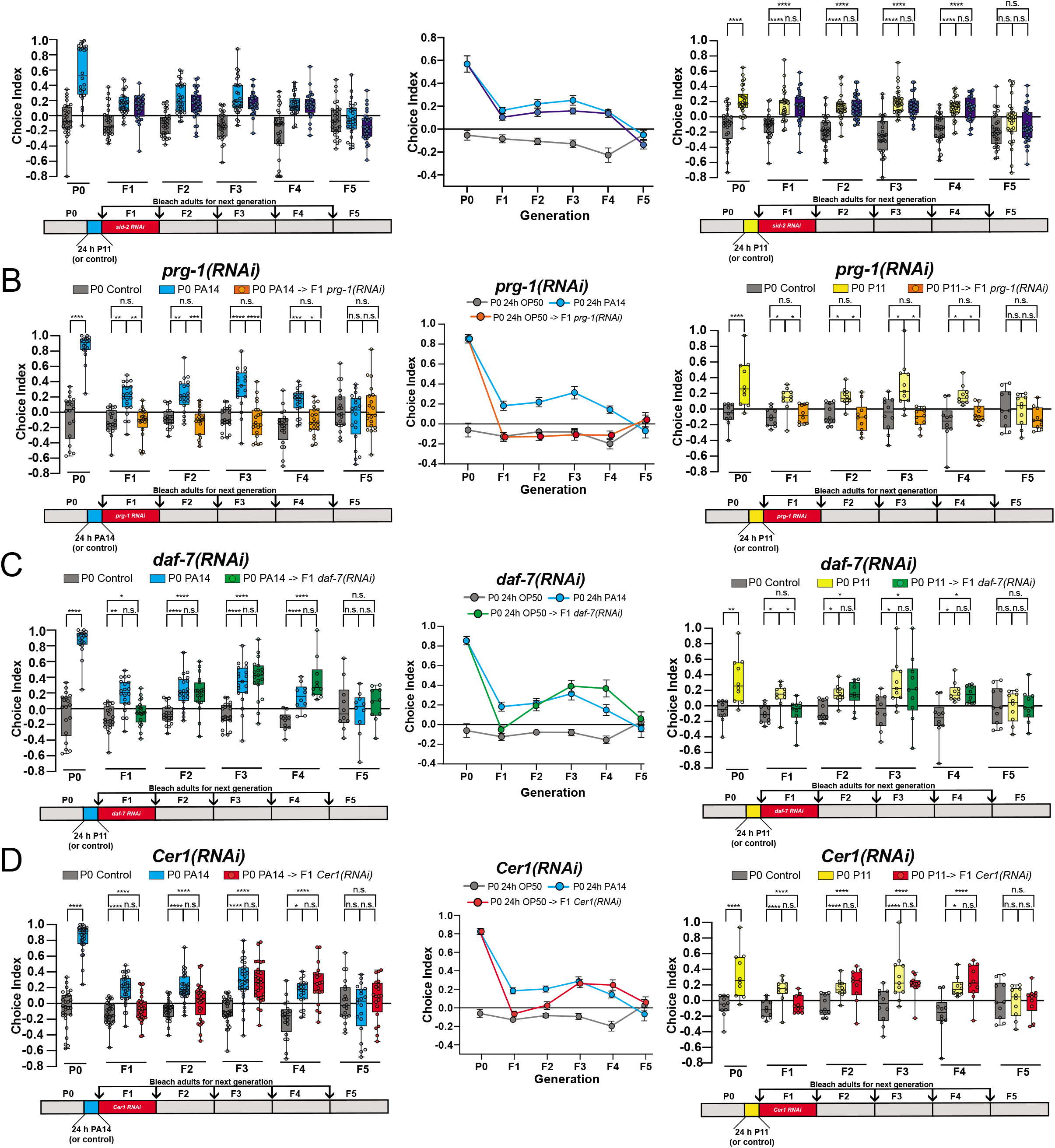
Transgenerational behavior effects of F2 generation-RNAi knockdown of *sid-2, prg-1, daf-7*, or *Cer1*. Wild-type mothers were trained with control, PA14 (left panels and line graphs) or P11-expressing *E. coli* (right panels). F1 progeny were then treated with either *sid-2* (B), *prg-1* (C), *daf-7* (D), or *Cer1* (E) RNAi. Subsequent generations of progeny were maintained on normal food and examined for PA14 avoidance behavior. Each dot represents an individual choice assay plate (average of 115 worms per plate) from all replicates. At least 3 biological replicates for all experiments. Box plots: center line, median; box range, 25-75th percentiles; whiskers denote minimum-maximum values. One-way (A-D) analysis of variance (ANOVA), Tukey’s multiple comparison test. * *P* < 0.05, \*\**P* ≤ 0.01, \*\*\**P* ≤ 0.001, \*\*\*\**P* < 0.0001, NS, not significant. Supplementary Table 1 for exact sample sizes (*n*) and *P* values.

**Supplemental Figure 6.**
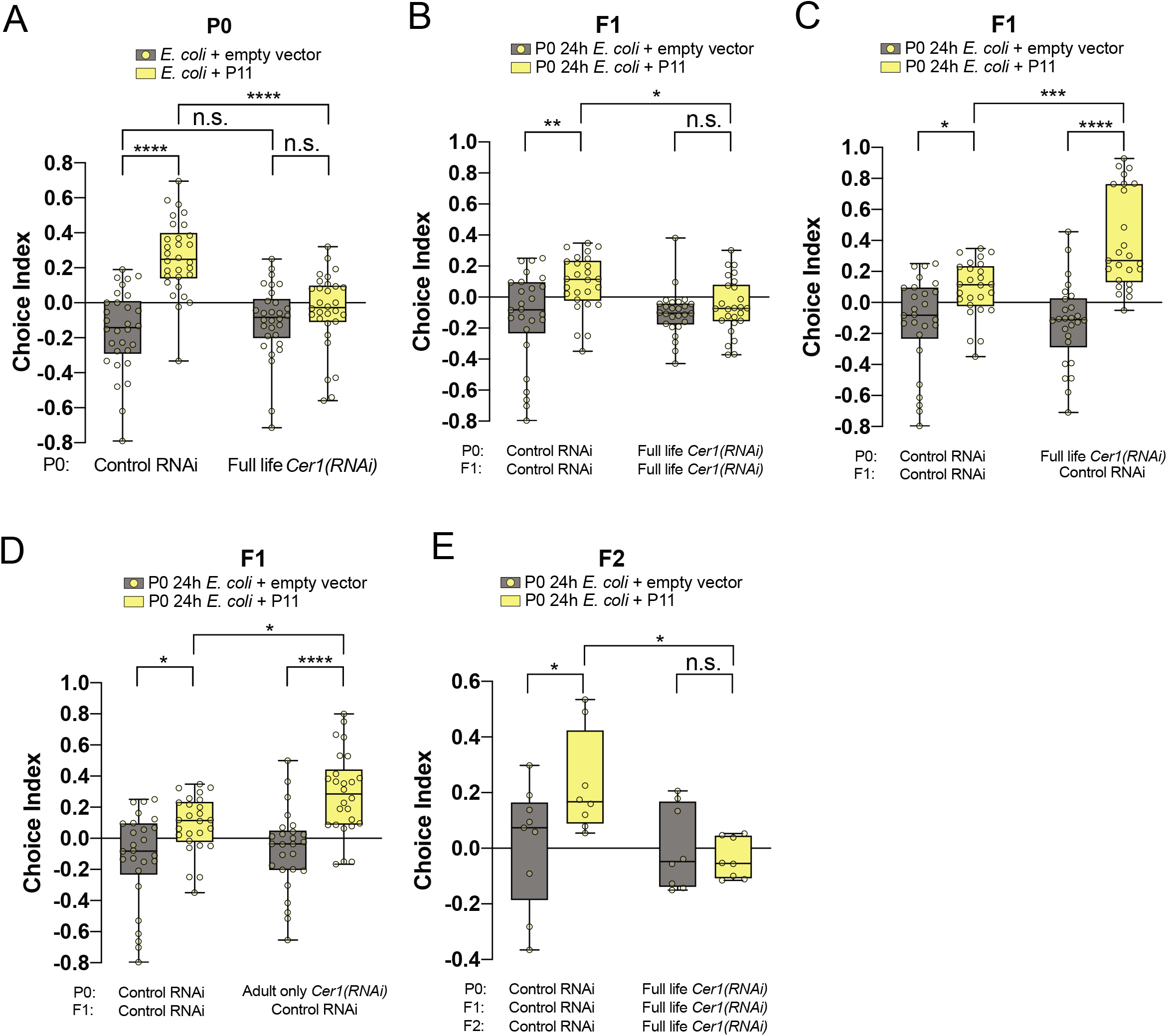
Adult-only RNAi knockdown of *Cer1*. A, Naïve mothers were treated from egg with *Cer1* or control RNAi. At the L4 stage, worms were trained on control or P11-expressing *E. coli* and tested for PA14 avoidance behavior. B, Progeny obtained from the trained mothers in (A) continued to be treated with whole-life control or *Cer1* RNAi. C, Progeny obtained from the trained mothers in (A) were treated only with control RNAi from egg to adulthood, then tested for PA14 avoidance behavior. D, Progeny obtained from the trained mothers in (Figure 4F) were treated only with control RNAi from egg to adulthood, then tested for PA14 avoidance behavior. E, The F2 grandprogeny from (A) and (B) continued to be treated with whole-life control or *Cer1* RNAi before PA14 avoidance behavior testing in adulthood. Each dot represents an individual choice assay plate (average of 115 worms per plate) from all replicates. At least 3 biological replicates for all experiments. Box plots: center line, median; box range, 25-75th percentiles; whiskers denote minimum-maximum values. Two-way (A-E) analysis of variance (ANOVA), Tukey’s multiple comparison test. * *P* < 0.05, \*\**P* ≤ 0.01, \*\*\**P* ≤ 0.001, \*\*\*\**P* < 0.0001, NS, not significant. Supplementary Table 1 for exact sample sizes (*n*) and *P* values.

**Supplemental Figure 7.**
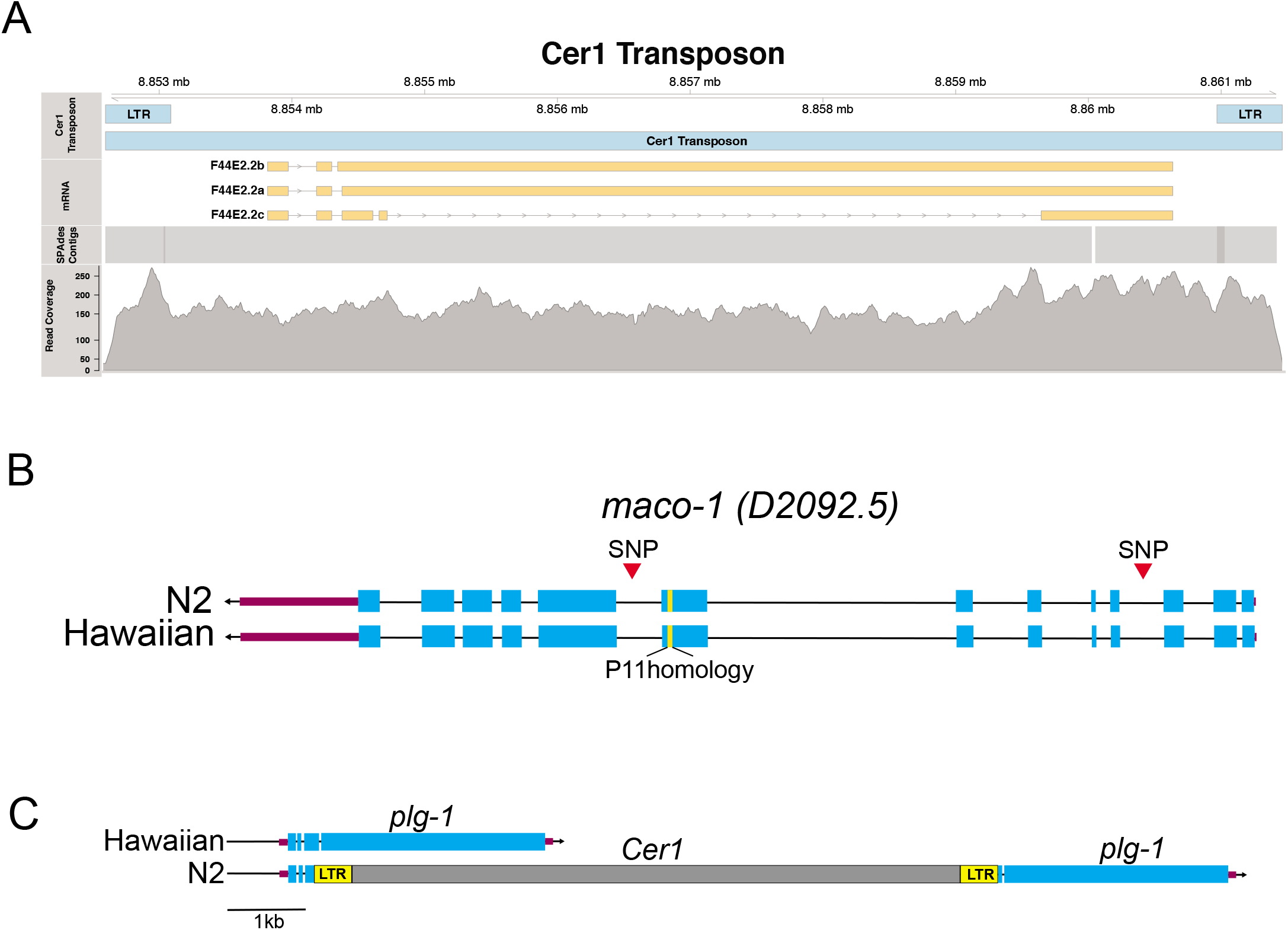
Genomic analysis of *Cer1, maco-1*, and *plg-1* loci. A, Full-length *Cer1* is present in JU1580, but not in the *plg-1* locus. B, The P11 mRNA target *maco-1* is intact in Hawaiian worms. C, Cer1 is inserted in the plg-1 locus of N2, but not Hawaiian.

**Supplemental Table 1. Statistics Reporting.**

Statistics were generated using Prism 8. Detailed results from all figures are provided.

## Materials & Methods

### C. elegans and bacterial strains cultivation

Worm strains were provided by the *C. elegans* Genetics Center (CGC): N2 (wild type), FK181, CB4856 (Hawaiian), JU1580, KR314, DH424, MY2, JU363, JU323, ED3077, ED3040, ED3054, ED3073 and, CB4037, and VC40895 (*gk870313*) (*Cer1* mutant). VC40895 was outcrossed 6 times to generate CQ655. CQ655 was used for all the *Cer1* mutant experiments reported in this paper.

Bacterial strains: *P. aeruginosa* PA14 was a gift from Z. Gitai. *P. fluorescens* 15 (Pf15) was a gift from M. Donia, OP50 was provided by the CGC, and *Serratia marcescens* (ATCC 274) was provided by the ATCC. Control (*L4440*), *Cer1, daf-7*, and *prg-1* RNAi clones were obtained from the Ahringer library and sequenced verified before use. *E. coli* expressing P11 was made as previously described (Kaletsky et al., 2020b).

General worm maintenance: Worm strains were maintained at 20°C on High Growth Media (HG) plates (3 g/L NaCl, 20 g/L Bacto-peptone, 30 g/L Bacto-agar in distilled water, with 4 mL/L cholesterol (5 mg/mL in ethanol), 1 mL/L 1M CaCl2, 1 mL/L 1M MgSO4, and 25 mL/L 1M potassium phosphate buffer (pH 6.0) added to molten agar after autoclaving) on *E. coli* OP50 using standard methods.

RNAi worm maintenance: For all experiments using control or *Cer1 RNAi* treated worms had been cultured on HG plates (supplemented with 1 mL/L 1M IPTG, and 1 mL/L 100 mg/mL carbenicillin) for at least three generations, never starving worms.

General bacterial cultivation: OP50 and *P. aeruginosa* PA14 were cultured overnight in autoclaved and cooled Luria Broth (10 g/L tryptone, 5 g/L yeast extract, 10 g/L NaCl in distilled water) shaking (250 rpm) at 37°C. *E. coli* strains expressing PA14 small RNAs were cultured overnight shaking (250 rpm) at 37°C in Luria Broth supplemented with 0.02% arabinose w/v and 100 mg/mL carbenicillin. *E. coli* RNAi strains were cultured overnight shaking (250 rpm) at 37°C in Luria Broth supplemented with filter sterilized 12.5 mg/mL tetracycline and 100 mg/mL carbenicillin.

### Training plate/worm preparation

Worm preparation: Eggs from young adult hermaphrodites were obtained by bleaching and subsequently placed onto HG plates seeded with *E. coli* OP50 or HG RNAi plates seeded with RNAi and incubated at 20°C for 2 days. Synchronized L4 worms were used in all training experiments.

Bacteria lawn training plate preparation: Overnight cultures of bacteria (prepared as described above) were diluted in LB to an Optical Density (OD_600_) = 1 and used to fully cover Nematode Growth Media (NGM) ((3 g/L NaCl, 2.5 g/L Bacto-peptone, 17 g/L Bacto-agar in distilled water, with 1 mL/L cholesterol (5 mg/mL in ethanol), 1 mL/L 1M CaCl2, 1 mL/L 1M MgSO4, and 25 mL/L 1M potassium phosphate buffer (pH 6.0) added to molten agar after autoclaving) plates. For preparation of *E. coli* expressing PA14 P11 RNA, bacteria were seeded on NGM plates supplemented with 0.02% arabinose and 100 mg/mL carbenicillin. All plates were incubated for 2 days at 25°C unless specified otherwise (in separate incubators for control/pathogen seeded plates). On the day of training (i.e., 2 days post bleaching), plates were left to cool on a benchtop for 1 hr to equilibrate to room temperature before the addition of worms. Additionally, for *E. coli* strains expressing PA14 small RNAs, 200 mL of 0.01% arabinose was spotted onto seeded training plates 1 hr prior to use.

Small RNA training plate preparation: 200 μL of OP50 was spotted in the center of a 10 cm NGM. Plates were incubated at 25°C for 2 days. 100 μg of small RNA was placed directly onto OP50 spots and left to completely dry at room temperature (~ 1 hr) before use on day of experiment for worm training.

### Worm preparation for training

Synchronized L4 worms were washed off plates using M9 and left to pellet on the bench top for approximately 5 minutes. 5 mL of worms were placed onto small RNA-spotted training plates, while 10 mL or 40 mL of worms were plated onto OP50 or *E. coli* expressing PA14 small RNAs, or pathogen-seeded training plates, respectively. Worms were incubated on training plates at 20°C in separate containers for 24 hr. After 24 hr, worms were washed off plates using M9 and washed an additional 3 times to remove excess bacteria. Worms were tested in an aversive learning assay described below.

### Aversive learning assay

Overnight bacterial cultures were diluted in LB to an Optical Density (OD_600_) = 1, and 25 mL of each bacterial suspension was spotted onto one side of a 60 mm NGM plate and incubated for 2 days at 25°C. After 2 days assay plates were left at room temperature for 1 h before use. Immediately before use, 1 mL of 1M sodium azide was spotted onto each respective bacteria spot to be used as a paralyzing agent during choice assay. To start the assay (modified from (Zhang et al., 2005)), worms were washed off training plates in M9 allowed to pellet by gravity, and washed 2 additional times in M9. 5 mL of worms were spotted at the bottom of the assay plate, using a wide orifice tip, midway between the bacterial lawns. Aversive learning assays were incubated at room temperature for 1 hr before manually counting the number of worms on each lawn. Plating a large number of worms (>200) on choice assay plates was avoided, since excess worms clump at bacterial spots making it difficult to distinguish animals, and high densities of worms can alter behavior.

In experiments in which each generation was treated with RNAi: Animals were washed off plates with M9 at Day 1 of adulthood. A subset of the pooled animals was subjected to an aversive learning assay, while the remaining worms were bleached to obtain eggs, which were then placed onto HG or HG RNAi plates and left at 20°C for 3 days before the next generation was tested.

### Statistical analysis of choice assay data

Populations of worms were raised together under identical conditions and were randomly distributed into treatment conditions. Trained worms were pooled and randomly chosen for choice assays. For all choice assays, each dot represents an individual choice assay plate (about 10–300 worms per plate) with all data shown from at least 3 independent replicates (Supplementary Table 1). Plates were excluded that contained less than 10 total worms per plate. The box extends from the 25th to the 75th percentile, with whiskers from the minimum to the maximum values. All figures in the Article and Supplementary Information pooled data from independent experiments. Statistics were generated using Prism 8. Counting of worms on choice assay plates was performed blind with respect to worm genotype and training condition.

### Preparation of bacteria for RNA isolation

Bacteria for RNA collection were prepared as described for training plates (i.e. 2 days on plates at 25°C). Bacterial lawns were collected from the surface of NGM plates using a cell scraper. Briefly, 1 mL of M9 buffer was applied to the surface of the bacterial lawn, and the bacterial suspension following scraping was transferred to a 15 mL conical tube. PA14, from 10 plates or OP50 from 15 plates was pooled in each tube and pelleted at 5,000 x g for 10 minutes at 4°C. The supernatant was discarded and the pellet was resuspended in 1 mL of Trizol LS for every 100 μL of bacterial pellet recovered. The pellet was resuspended by vortexing and subsequently frozen at −80°C until RNA isolation.

### Bacteria RNA isolation

To isolate RNA from bacterial pellets, Trizol lysates were incubated at 65°C for 10 min with occasional vortexing. Debris was pelleted at 7000 x g for 5 min at 4°C. The supernatant was transferred to new tubes containing 1/5 the volume of chloroform. Samples were mixed thoroughly by inverting and centrifuged at 12000 x g for 10 min at 4°C. The aqueous phase was used at input for RNA purification using the mirVana miRNA isolation kit according to the manufacturer’s instructions small RNA (<200 nt) isolation. Purified RNA was used immediately or frozen at −80°C until further use as previously described (Kaletsky et al., 2020a).

### C. elegans total RNA isolation

F2 worms from trained grandmothers were washed off of plates using M9. Three additional M9 washed were performed to remove excess bacteria, and the supernatant was discarded. 1 mL of Trizol LS was added per 100_ μL of worm pellet. Worms were lysed in Trizol by incubation at 65°C for 10 min with occasional vortexing. RNA was extracted with chloroform, and the aqueous phase was used as input for RNA purification using the mirVana miRNA isolation kit according to the manufacturer’s instructions for total RNA. Approximately 100μg of total RNA from either control or P11 grandmother-trained F2 worms was used per training plate. This amount of RNA was chosen as it correlates to the same input of worms used for training with worm lysate (see Preparation of Worm Lysates). Purified RNA was used immediately by dropping RNA onto preseeded spots of OP50 on NGM plates. Plates were allowed to air dry before the addition of naïve worms for training. Worms were trained on RNA-seeded plates for 24 h at 20°C and subsequently tested for PA14 aversive learning using a standard choice assay.

### Analysis of JU1580 genomic sequences

Fastq files from SRA (accession numbers SRR9322509, SRR9322510, SRR9322511, SRR9322512, SRR9322514) were uploaded to Galaxy (Afgan et al., 2018) for analysis. De novo assembly of Illumina reads was performed using SPAdes (Bankevich et al., 2012) (Galaxy wrapper version 3.12.0), and contigs were aligned to the *C. elegans* N2 strain genome (WBcel235) using minimap2 (Li, 2018). For structural variant detection, alignment of raw fastq reads to *C. elegans* was performed using BWA (Li and Durbin, 2009), followed by analysis using Lumpy (Layer et al., 2014).

### Preparation of worm lysates

Day 1 F2 progeny from control or P11-trained grandmothers were collected from plates and washed 3 times in M9. The worm pellet was washed with DPBS, and the pellet was resuspended in DPBS. Worms were homogenized using an all-glass Dounce tissue grinder (Kimble # 885300-0002), and homogenization was monitored using a microscope. Different worm lysates within an experiment were normalized to the starting amount of worms. For training naïve worms with lysates from F2 animals, the normalized lysate was diluted 1:3 with DPBS, such that 400 uL of lysate was obtained for every 100 ul of starting worm pellet. 150 ul of lysate was immediately pipetted directly onto the bacterial spot of 10 cm NGM plate (seeded with 200 ul of an OP50 spot in the center of the plate, 2 days prior to the experiment). Worm lysates were allowed to air dry, and plates with lysates were monitored to ensure no worms were alive following homogenization. Naïve Day 1 worms were then transferred to lysate-seeded plates for 24h of training at 20°C, followed by testing for learned avoidance using the standard OP50 v. PA14 choice assay.

### Cer1-enriched fraction isolation

Homogenates were prepared as described (Preparation of worm lysates) and cleared from debris by a 750 x g centrifugation at 4° C for 5 minutes. Homogenization and clearing steps were repeated twice. The homogenates were then passed twice through a 0.22 um filter. For each sample, the homogenate protein concentration was measured using Quant-iT Protein Assay Kit (Invitrogen #Q33211). Per experiment, if needed, the homogenates were diluted in DPBS in order to load similar concentrations. From each sample, a small aliquot was kept as a “load” sample, and 830 uL was layered on top of an Iodixanol gradient. For each gradient-5%, 11%, 17%, 24% and 30% Iodixanol solutions were made by mixing solution A (0.1 M NaCl, 0.5 mM EDTA, 50 mM Tris HCl, pH 7.4) with solution B [50% Iodixanol solution (OptiPrep, Sigma #D1556), 0.5 mM EDTA, and 50 mM Tris HCl, pH 7.4]. The gradient was made in a 5 mL, Open-Top Thinwall Ultra-Clear Tube (Beckman Coulter #344057) from equal volumes (830 uL) of each Iodixanol solution that were allowed to diffuse by an overnight incubation at 4°C. Samples were then centrifuged at 112,000 x g (4° C) for 2 hours, using SW55 Ti Swinging-Bucket Rotor (Beckman Coulter). Six fractions of equal volumes were collected. *Cer1*-enriched fraction (fraction 6), as well as fraction 3, were diluted in DPBS and centrifuged at 335,000 x g (4° C) for 30 minutes. Each pellet was then resuspended in DPBS and used for Western blots, naïve worm training, or electron microscopy. For each experiment, the enrichment of *Cer1* in fraction 6 was verified by western blot. For fractions treated with RNase, 1:1000 RNaseA (omega BIO-TEK #AC117) was added following resuspension after the final spin, and samples were incubated for 15 minutes at room temperature. For behavior experiments with RNase-treated samples, the reaction was terminated by adding RNase inhibitor (Invitrogen #AM2696, 1 unit final).

#### Western blot

For western blot analysis, samples were mixed with 10X Bolt Sample Reducing Agent (Invitrogen #B0009) and 4X Bolt LDS Sample Buffer (Invitrogen #B0007). Samples were then heated at 70°C for 10 minutes before loading on a gradient-PAGE (4 % −12 %) Bis-Tris gel (Invitrogen #NW04125BOX). After their separation, samples were transferred to a PVDF membrane (Millipore #IPVH00010) and blocked with 5 % milk in TBST (10X TBST: 200 mM Tris-HCl, pH 7.5, 1.5 M NaCl, 1% Tween20). Membranes were incubated with one of the following primary antibodies: anti-Cer1 GAG, 1:150 dilution), rabbit polyclonal anti-Histone H3 (Abcam #ab1791, 1:1000 dilution), mouse monoclonal anti-ATP5A (Abcam #ab14748, 1:1000 dilution), mouse monoclonal anti-Hsp90 (Abcam #ab13492, 1:1000 dilution). After washing with TBST, membranes were incubated with the corresponding fluorescent secondary antibody (either goat anti-rabbit IgG, Invitrogen #A32732 or goat anti-mouse IgG-Invitrogen #A28175). Membranes were then washed with TBST and imaged on (ODYSSEY CLx).

### Imaging and fluorescence quantification

All images were taken on a Nikon Eclipse Ti microscope. Differential inference contrast (DIC) images of whole worms following OP50, or PA14 lawn or small RNA training, were imaged at 20×. *Z*-stack multi-channel (DIC and GFP) of day-1 adult GFP-transgenic worms were imaged every 1 μm at 60× magnification; Maximum intensity projections and 3D reconstructions of head neurons were built with Nikon NIS-Elements. To quantify *daf-7p::gfp* levels, worms were prepared and treated as described in ‘Worm preparation for training’. Worms were mounted on agar pads and immobilized using 1 mM levamisole. GFP was imaged at 60× magnification and quantified using NIS-Elements software. Average pixel intensity was measured in each worm by drawing a Bezier outline of the neuron cell body for 2 ASI head neurons.

### Whole-mount immunofluorescence

Antibody staining of *C. elegans* gonads (Annette Chan and Barbara Meyer, Wormbook: Methods in Cell Biology). Day 1 hermaphrodites were suspended in M9 on a glass slide and gonads were dissected. Slides were freeze-cracked on dry-ice, fixed for 5 min in cold MeOH/5 min in EtOH, and washed 3x in PBST. Primary antibodies used: anti-*Cer1*-GAG (1:50), and anti-Histone H3 (Abcam, 1:200). Secondary antibodies used: goat anti-mouse AlexaFluor 488-labeled IgG (1:500), goat-anti rabbit AlexaFluor 555-labeled IgG (1:500). Both primary and secondary antibodies were incubated overnight at 4C. Images were taken with a Nikon Ti at 40x.

### Negative-stain Electron Microscopy

5 μl of samples were applied to glow-discharged grids (Electron Microscopy Sciences, CF400-Cu), washed once with ultrapure water, and stained with 0.75% uranyl formate. Images were collected with a Talos F200X Transmission Electron Microscope with CCD camera, at 200 keV at magnifications of either 14,000 (lower mag) or 36,000 (higher mag).

